# Endosomal maturation is controlled by the trimeric Bulli-Mon1-Ccz1 GEF7 complex and Rab5-GTPase activating protein GAPsec

**DOI:** 10.64898/2026.03.05.709801

**Authors:** Maren Janz, Maik Drechsler, Heiko Meyer, Vikeraman Sriram, Kim Michelle Simes, Elisa Frommhold, Nadia Füllbrunn, Lars Langemeyer, Christian Ungermann, Daniel Kümmel, Achim Paululat

## Abstract

The endolysosomal system is crucial for the degradation of cellular waste in the lysosomal lumen. Within this pathway, endosomes mature prior to their fusion with lysosomes. This process relies on the sequential action of the CORVET and HOPS tethering complexes, guided by Rab5 and Rab7 GTPases, respectively. CORVET acts on early endosomes (EEs), transitioning to HOPS on maturing late endosomes/multivesicular bodies (LEs/MVBs) for lysosomal fusion. This process is finely tuned by the Rab activating guanine nucleotide exchange factor (GEF) and the inactivating GTPase activating protein (GAP). The BuMC1 GEF complex (Bulli-Mon1-Ccz1) uniquely activates Rab7 in metazoans and interacts with Rab5, which stimulates its activity. Here, we identified GAPsec as a novel GAP with activity for Rab5 required for endosomal maturation in fruit fly nephrocytes. Inactivation of GAPsec results in enlarged, dysfunctional endosomes that are unable to reach lysosomes for degradation. Our study highlights the importance of coordinated Rab regulation for efficient endosomal trafficking.

## INTRODUCTION

The maturation of early endosomes into late endosomes/MVBs and fusion with lysosomes in all eukaryotes is highly dependent on the hexameric tethering complexes CORVET and HOPS. In *Drosophila*, a tetrameric miniCORVET acts as an endosomal tether (Lőrincz et al., 2019; Lőrincz et al., 2016). CORVET and its associated GTPase Rab5 act on early endosomes and are replaced upon endosomal maturation by HOPS and its associated GTPase Rab7 on MVBs/LEs. HOPS promotes the fusion of LEs with lysosomes and is thus critical for the degradation of unwanted compounds in the cell (Balderhaar et al., 2013; Balderhaar and Ungermann, 2013; Poteryaev et al., 2010). It has also been shown that, besides Rab7, other small GTPases are also required for endosome maturation. Rab2 and Arl8 have both been implicated in endosome-lysosome fusion and are thus crucial for the delivery of functional lysosomes in nephrocytes (Boda et al., 2019; Lőrincz et al., 2017; Lund et al., 2018).

The yeast, fly, and mammalian CORVET complex binds directly to Rab5 at the membrane of early endosomes, promoting their maturation. It is assumed that Rab5 binds to the EE and recruits the CORVET complex towards the endosomal membrane with the assistance of additional effector proteins such as Rabaptin-5, Rabex-5, and others. Later, HOPS is recruited via Rab7 on the late endosome, thus mediating membrane tethering between the late endosome and lysosome. However, Rab7 has additional cellular functions; for example, it interacts with the retromer complex to control endosomal positioning (Balderhaar et al., 2010; Bonifacino and Neefjes, 2017; Jimenez-Orgaz et al., 2018; Liu et al., 2012; Purushothaman et al., 2017; Rojas et al., 2008).

Rab protein activation is regulated by the exchange of bound GDP for the more abundant GTP, mediated by specific GEF proteins (guanine nucleotide exchange factors). GAPs (GTPase-activating proteins) promote the hydrolysis of the γ-phosphate, resulting in the inactivation of Rab, followed by its subsequent membrane extraction via the chaperone-like GDP dissociation inhibitor (GDI). During endosomal maturation, the highly coordinated activity of GEFs and GAPs is essential for maintaining correct and timely coordinated cellular trafficking. In yeast, the Mon1-Ccz1 heterodimer acts as a major Rab7-GEF (Nordmann et al., 2010). In metazoans, the Mon1-Ccz1 GEF complex has a third subunit, which is required for GEF function *in vivo* (Dehnen et al., 2020; Herrmann et al., 2023b; Vaites et al., 2018; Yong et al., 2023), though not for GEF activity *in vitro* (Borchers et al., 2023; Langemeyer et al., 2020). We identified this third subunit in *Drosophila,* named Bulli, and thus refer to the trimeric complex as BuMC1 (Dehnen et al., 2020). We have previously shown that Bulli, the unique metazoan component of the complex, increases GEF activity towards Rab7 by approximately 20% *in vitro*, which is considered highly important in a physiological context (Dehnen et al., 2020). A lack of Bulli in *Drosophila* nephrocytes results in multiple cellular phenotypes associated with the disruption of endomembrane trafficking pathways. A particularly prominent phenotype is the appearance of giant vesicles, which we assume to be endosome-like compartments resulting from misregulated vesicle fusion. These “late endosome”-like vesicles fail to become Rab7-positive. Consequently, they do not acidify and thus fail to fuse with lysosomes (Dehnen et al., 2020).

Moreover, we recently showed that the Rab7-GEF activity of the BuMC1 complex is stimulated by Rab5 in a reconstituted system (Borchers et al., 2023). Direct interaction with Rab5 occurs within a conserved region of the LONGIN domain 2 (LD2) of Mon1 (Borchers et al, 2023). Structural analysis of the BuMC1 complex suggests that Bulli does not affect the catalytic function of the core complex by interacting with the active sites but rather represents a docking platform (Herrmann et al, 2023b). Indeed, the membrane binding of the metazoan BuMC1 complex requires Bulli (Wilmes et al., 2025). Our simulation studies revealed that basic residues in the ß-propeller of Bulli mediate electrostatic interaction with membranes. Moreover, mutations in the ß-propeller of Bulli as a membrane interface impaired targeting of Bulli to membranes and thus BuMC1 function in transgenic fly lines (Wilmes et al., 2025). The studies show that the trimeric BuMC1 complex requires dual interactions during endosomal maturation: one with Rab5 via Mon1 and another with membranes via the Bulli subunit. We do not yet know whether Rab5 binding is essential for the subsequent activation of Rab7, though BuMC1 is required for the recruitment and activation of Rab7 at the LE.

In the present work, we demonstrate that the loss of Mon1 or Ccz1, the two other constituents of the BuMC1 complex, causes cellular phenotypes in nephrocytes very similar to the loss of Bulli. This indicates that all three components of the Rab7-GEF complex are equally critical for the transition of EE to LE and for the proper activation of Rab7. We also investigated the role of the BuMC1 complex in endosome formation by interfering with the maturation process via the expression of constitutively active and dominant-negative mutants of Rab5 and Rab7. We found that inactivation of Rab5 at early endosomes is crucial for membrane release and the continuing endosomal maturation, and this process requires the BuMC1 complex and activation of Rab7. In addition, we asked for the role of GAPs in endosomal maturation in nephrocytes and performed an unbiased RNAi-mediated knockdown screen. 24 of the 25 annotated GAPs in the *Drosophila* genome were specifically down-regulated in nephrocytes. Cellular consequences were microscopically analysed by Rab5 and Rab7 staining, which led to the identification of two new candidate Rab GAPs, GAPsec and Tbc1d22, that are required for endosomal maturation. *In vitro* assays showed GAP activity of GAPsec towards Rab5, suggesting a role for GAPsec as Rab5-GAP in *Drosophila* nephrocytes.

## RESULTS

### Bulli/Mon1/Ccz1 (BuMC1) acts as a complex in nephrocyte endocytosis

Loss-of-function mutations in Bulli cause dramatic alterations in the morphology of endo-lysosomal structures, as well as a reduction of the endocytic capacity in pericardial nephrocytes in *Drosophila* (Dehnen et al., 2020). *bulli* mutant nephrocytes exhibit three major phenotypes – the formation of enlarged vesicles, the loss of efficient Rab7 recruitment required for endosome maturation, and the intravesicular accumulation of Rab5 in endosomal carriers (Dehnen et al., 2020). While the impairment of Rab7 recruitment can be explained by the Rab7-GEF function of the BuMC1 complex, the accumulation of Rab5 has not been addressed in detail thus far. To investigate whether the deposition of Rab5 in enlarged endosomes is solely due to the absence of Bulli or whether it constitutes a dysfunction of the entire trimeric BuMC1 complex, we analysed Rab5 and Rab7 localisation and intracellular distribution in *mon1*^mut4^ and *ccz1*^d113^ mutants by immunostaining (Figure 1). Similar to *bulli*^6-61^ mutants, we observed that the lack of either Mon1 or Ccz1 results in intravesicular Rab5 deposits throughout the cells (Figure 1B-D). In addition, we investigated the ultrastructure of nephrocytes from *mon1*^mut4^ and *ccz1*^d113^ loss-of-function mutants in comparison to the ultrastructural phenotypes of *bulli*^6-61^ mutants (Figure 2) (Dehnen et al., 2020). All three mutants exhibited comparable, but not fully identical phenotypes, including the appearance of enlarged endosomal vesicles with internalised Rab5 deposits.

**Figure 1:**
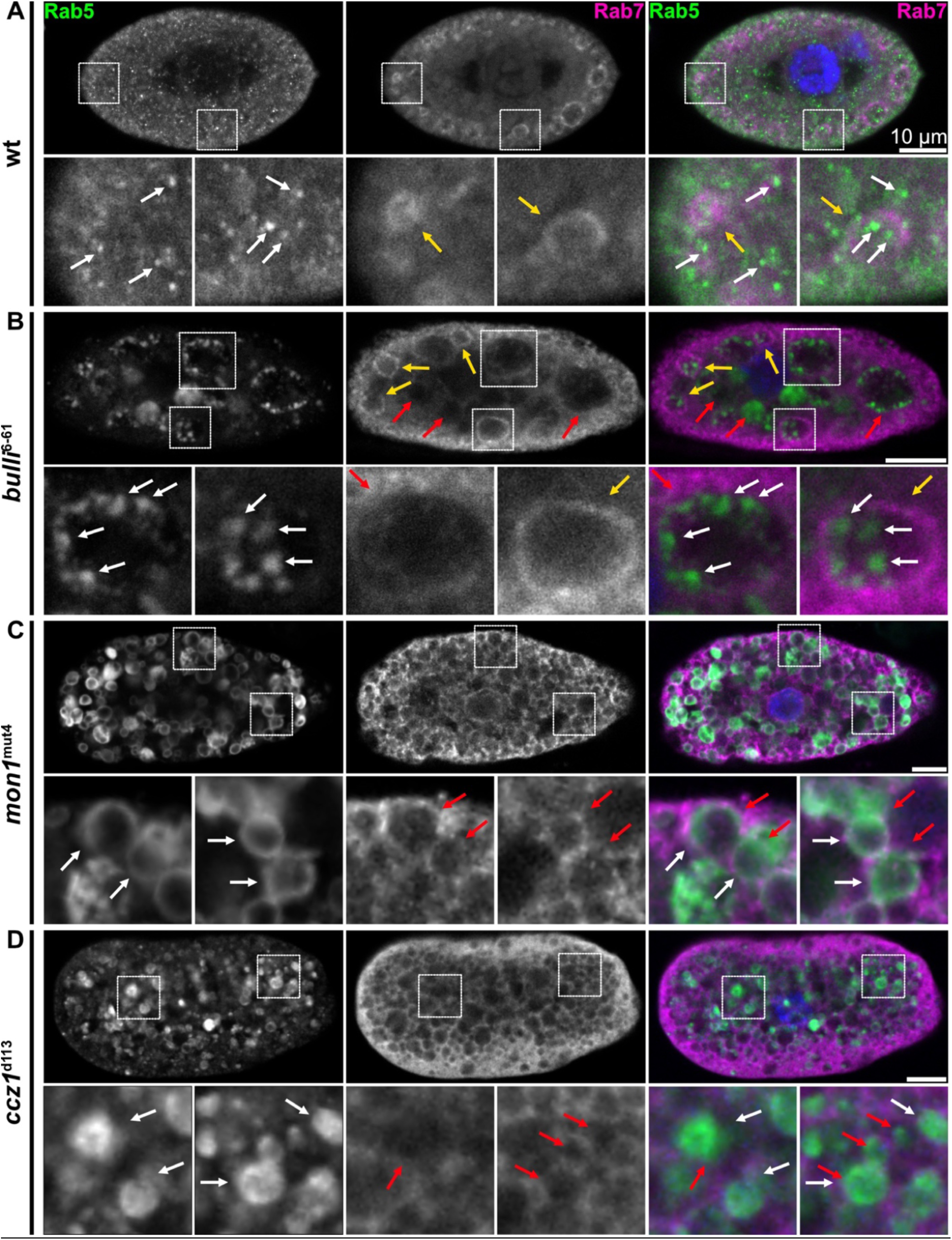
Effect of loss-of-function mutations in *bulli*, *mon1* and *ccz1* on endolysosomal biogenesis. Pericardial nephrocytes from dissected 3^rd^ instar larvae of control flies (A) and *bulli*^6-61^ (B), *mon1*^mut4^ (C) and *ccz1*^d113^ (D) mutants were simultaneously stained for Rab5 and Rab7. Single channels, merged images and close-ups were shown. White arrows illustrate Rab5 signals, yellow arrows highlight Rab7-positive vesicles, and red arrows point to Rab7-negative enlarged vesicles. Scale bar indicates 10 μm.

**Figure 2:**
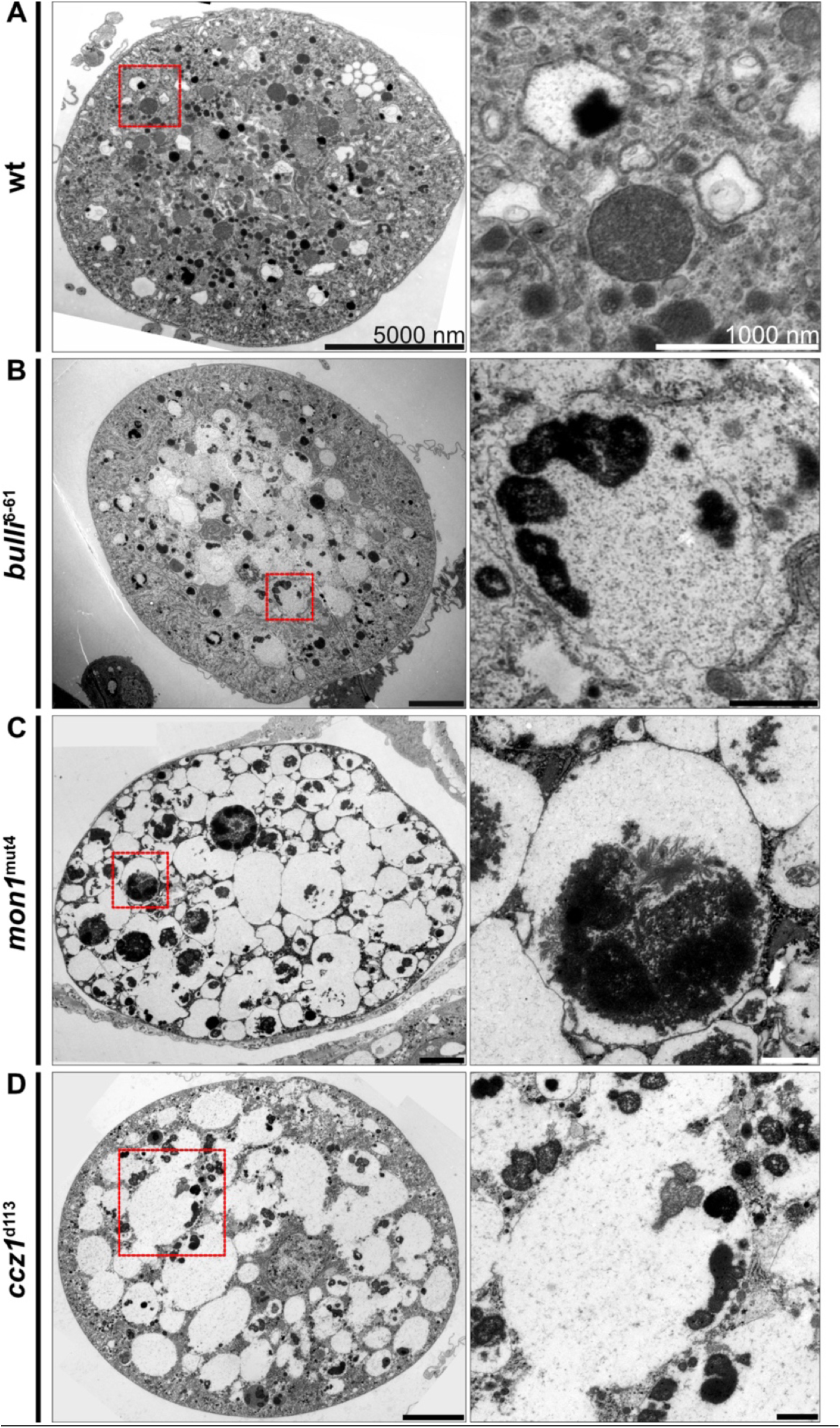
Ultrastructural defects in nephrocytes in *bulli*, *mon1*, and *ccz1* mutants. Ultrastructural analysis of nephrocytes from dissected 3rd instar larvae of control flies (A) and *bulli*^6-61^ (B), *mon1*^mut4^ (C) and *ccz1*^d113^ (D) mutants. In all analysed mutants, we found highly enlarged endosomal vesicles compared to those in control animals. The enlarged vesicles contain electron-dense aggregated material. We have previously demonstrated that this corresponds to membrane remnants with anchored Rab5 in *bulli*^6-61^ mutants (Dehnen et al., 2020).

Nephrocytes adapt by several ways to the specialised needs of scavenger cells: (i) they grow by polyploidism to cells of more than 100 µm diameter in size, thus offering enormous storage capacity (Kambysellis and Wheeler, 1972; Rizki, 1978). All nephrocytes serve for a life-time, they are not renewed. (ii) Endocytic and degradation pathway genes are all highly expressed in nephrocytes to ensure highest clearance and recycling activity (Meyer et al., 2024), (iii) a large array of membrane invaginations constitute a labyrinth channel system, increasing the endocytic active zone enormously (Das et al., 2008; Lehmacher et al., 2012; Miyaki et al., 2020; Psathaki et al., 2018; Psathaki and Paululat, 2022), and (iv) they harbour a 70 kDa exclusion size filtration barrier at the entries to the labyrinth channels, built by the overlaying ECM and by slit diaphragms, which represent specialised adherens junctions required to form the entry pores to the labyrinth channels (Carrasco-Rando et al., 2023; Gass et al., 2022; Helmstädter et al., 2017; Ivy et al., 2015; Lang et al., 2022; Weavers et al., 2009). Therefore, we analysed the labyrinth channel system of nephrocytes to check for effects of mutations in the BuMC1 complex (Figure 3).

**Figure 3:**
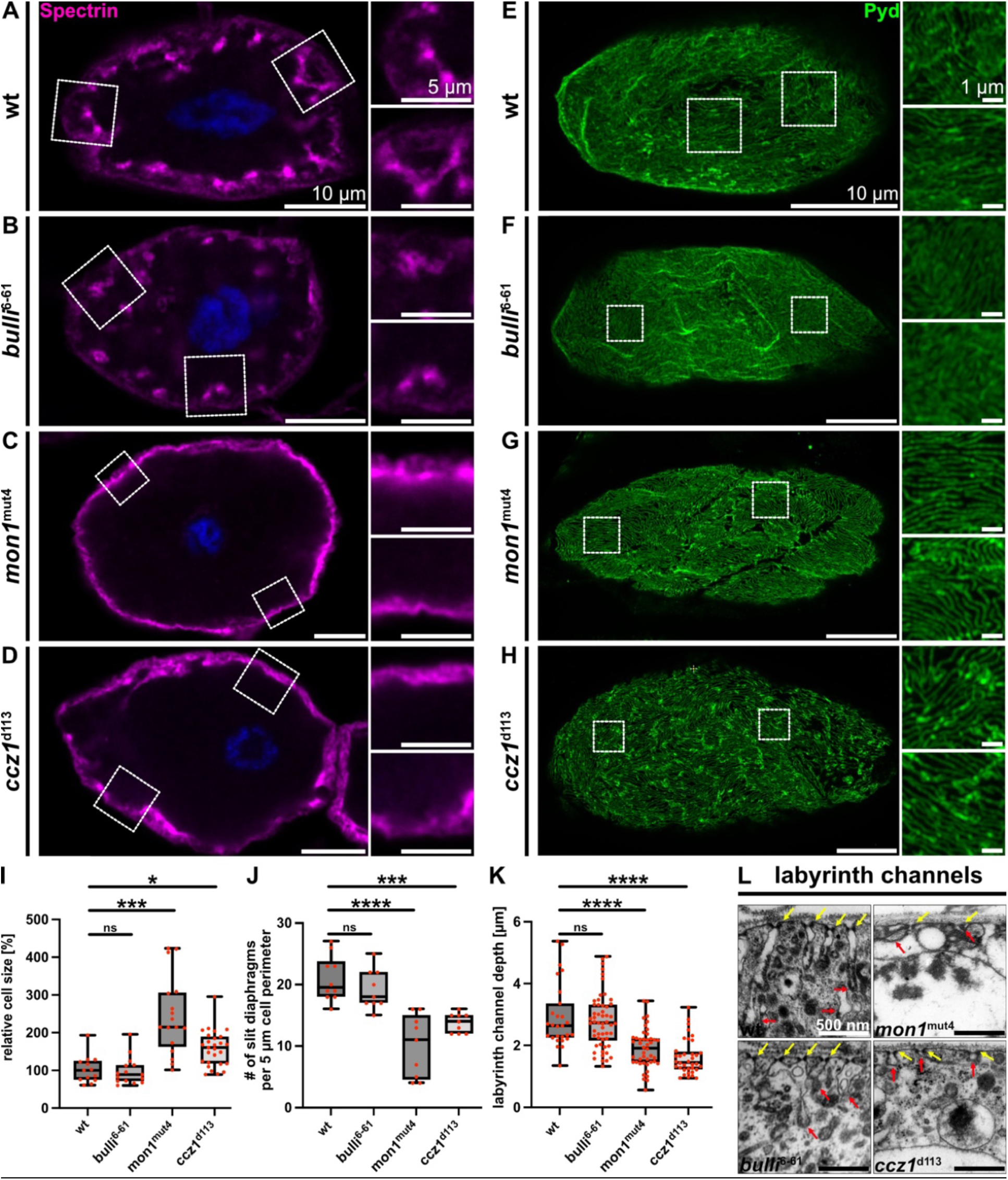
Spectrin and Pyd localisation in wildtype and in *bulli*, *mon1*, and *ccz1* mutant nephrocytes. Nephrocytes from dissected 3^rd^ instar larvae of wildtype (A, E), *bulli*^6-61^ (B, F), *mon1*^mut4^ (C, G) or *ccz1* ^d113^ mutants (D, H) were stained with spectrin or Pyd antibodies. Close-ups (indicated by dotted boxes in the overviews) are shown. Scale bars indicate 10 μm, 5 µm, and 1 µm, respectively. (I) Relative size of nephrocytes of the depicted genetic backgrounds. (J) Number of slit diaphragms (SDs) per 5 μm of cell perimeter, based on TEM images shown in Figure 2. (K) Quantification of the width of the cortical labyrinth channel system in wildtype and mutant nephrocytes, as evaluated by spectrin labelling. (L) Representative images of the labyrinth channel system in the depicted genetic backgrounds (TEM images).

Compared to wildtype, mutations in *mon1* and *ccz1* cause several deficits in the morphology of the labyrinth channel system. When stained for spectrin, a cytoskeletal protein that lines the intracellular side of the plasma membrane and that we used as a readout for the depth of labyrinth channel invagination (Zarnescu and Thomas, 1999), we found a significant decrease of the width of the cortical invagination zone in *mon1* and *ccz1* mutants compared to wildtype (Figure 3A-D, K). When stained for Polychaetoid/ZO-1 (Pyd), a marker for slit diaphragms (SDs) (Carrasco-Rando et al., 2019), we observed that SDs form in all mutant backgrounds but are spaced farther apart (Figure 3E-H, J). We quantified the size of nephrocytes in BuMC1 mutants relative to wildtype and found that the nephrocytes in *mon1* and *ccz1* mutants are bigger than control cells (Figure 3I). This might be caused by the uncontrolled fusion of endosomes and the occurrence of enlarged endosomes, which, as a consequence, expand the cell. Interestingly, the number of SDs measured alongside the cell perimeter was significantly reduced in *mon1* and *ccz1* mutants, but not in *bulli* mutants (Figure 3J). However, this effect may in part be caused by the concomitant increase in cellular volume that may move the SDs somewhat apart. This is also indicated in the Pyd-stainings (Figure 3E-H). Interestingly, using the measurement methods we applied here, we cannot detect any significant effects in the labyrinth channel system or in the formation or localisation of the slit diaphragms in *bulli* mutants, which differs from our findings for *mon1* and *ccz1*. However, our previous work has shown that the dimeric Mon-Ccz1 complex exhibits high GEF activity on its own, which increases by 40% when Bulli is added to the GEF assay (Dehnen et al., 2020). While Bulli, together with Mon1 and Ccz1, is essential for EE maturation, Bulli appears to play a minor role in maintaining the labyrinth channel system.

Taken together, Rab5 accumulation and malformation of the labyrinth channels do not constitute a *bulli*-specific defect in the endosomal pathway, but are caused by a dysfunction of the entire trimeric BuMC1-complex. Mutations in any of the three proteins of the trimeric BuMC1 complex result in a failure of Rab5 clearance from EEs, and Rab7 recruitment to endosomal membranes, indicating two distinct but connected roles of the BuMC1 complex in endosome maturation. Based on these data, we postulate that a dysfunctional trimeric Rab7GEF complex first prevents the inactivation and release of Rab5 from early endosomes and, second, causes a failure to recruit Rab7 towards the endosomal membrane, thus inhibiting maturation to LEs.

### Expression of Rab5^CA^ causes Rab5 internalisation into endosomes

To test whether a prolonged association of Rab5 with the membrane of early endosomes results in its intraluminal deposition within endosomes, we overexpressed a GTP-locked (constitutively active) version of Rab5 (Rab5^CA^; Q88L) in nephrocytes and investigated the subcellular distribution of Rab5 and Rab7 (Figure 4). Similar to mutations in the trimeric BuMC1 complex, overexpression of Rab5^CA^ induces the formation of enlarged vesicles, negative or positive for Rab7, that show an accumulation of intravesicular Rab5 protein or Rab7-positive vesicular structures that are wrapped around Rab5-positive membranes (Figure 4B). Importantly, the observed Rab5 accumulation mimics the phenotype seen in BuMC1 mutants, suggesting that these are aberrant endosomes. This further suggests that the absence of BuMC1 leads to misregulation of Rab5 activity. We also tested the consequences of expressing a GDP-locked (dominant negative) version of Rab5 (Rab5^DN^; S43N) in an otherwise wildtype background (Figure 4C). Under these conditions, the nephrocytes appear normal for endosomal maturation. When cells expressing Rab5^DN^ were stained for endogenous Rab5 and Rab7, both proteins localised to EE and LE, comparable to control cells. It is known that Rab5^DN^ remains largely cytosolic, bound to GDI-complexes, and fails to bind to EE membranes (Edler and Stein, 2019). Thus, as expected, we do not see an increased cytosolic Rab5 signal, likely due to the fast degradation of Rab5^DN^. However, Rab7 localises like wildtype in Rab5^DN^ flies (yellow arrows). Our results indicate that a prolonged membrane association of GTP-locked Rab5 mimics the Rab5 clustering phenotype observed in any of the BuMC1 mutants.

**Figure 4:**
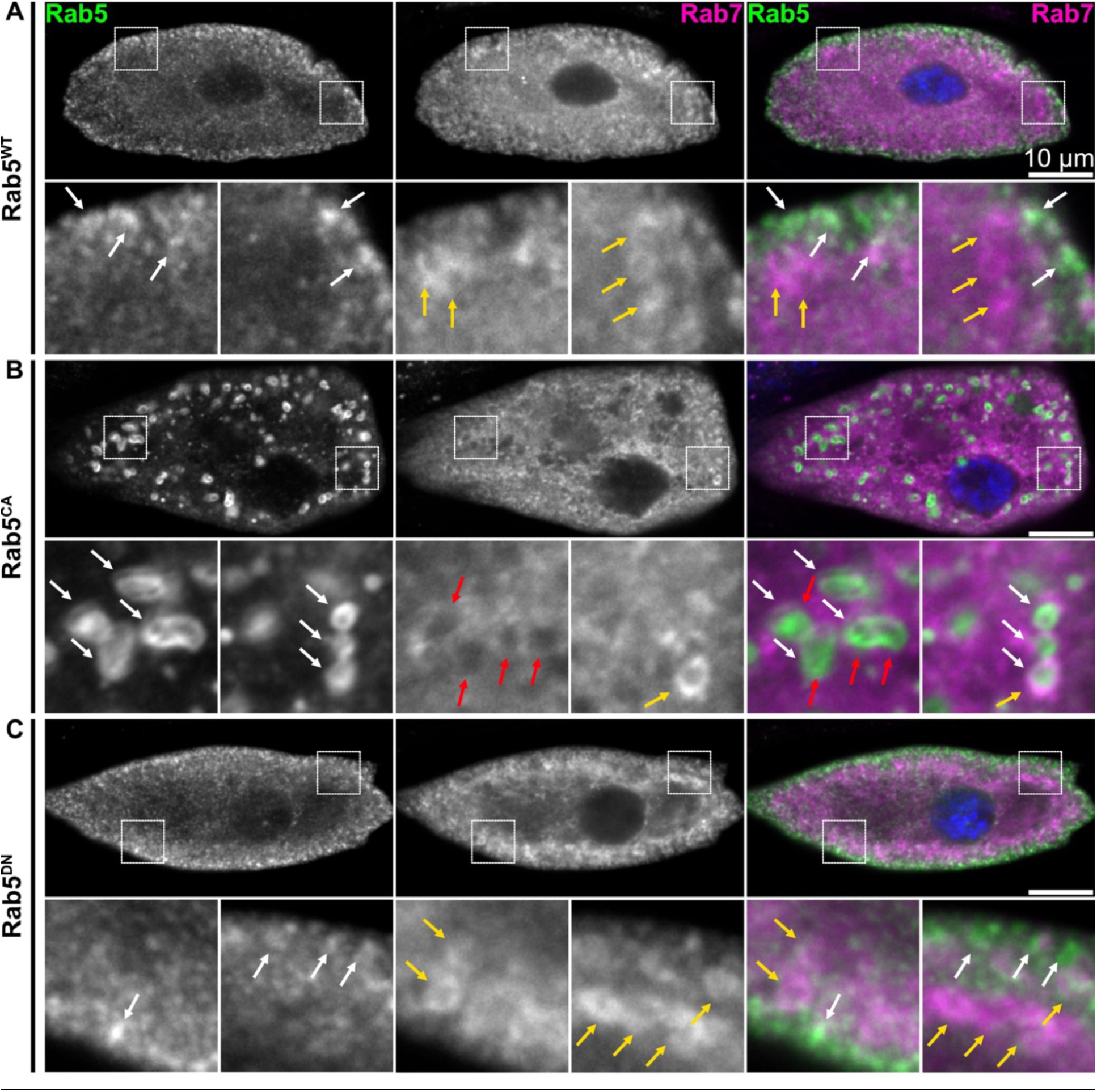
Expression of Rab5^ca^ causes Rab5 internalisation into endosomes. Nephrocytes from dissected 3^rd^ instar larvae of flies, in which either Rab5^WT^ (A), Rab5^CA^ (B)or Rab5^DN^ (C) was expressed on top of the endogenous Rab5, were stained with Rab5 and Rab7 antibodies. Expression of the three constructs, GFP.Rab5^WT^, YFP.Rab5^CA^ and YFP.Rab5^DN^ was driven by *hand*C-Gal4. Single channels, merged images and close-ups (indicated by dotted-lined boxes in the overviews) were shown. Enlarged endocytic vesicles were found in flies, in which Rab5^CA^ was expressed (white arrows). Scale bar indicates 10 μm.

### Activity of Rab7 impacts Rab5 clearance at endosomes

The maturation of early to late endosomes is driven by a Rab5-to-Rab7 switch on the endosomal membrane (Borchers et al., 2021). While Rab5 becomes inactivated by GTP hydrolysis, assisted by one or more GTPase-activating enzymes (RabGAPs), and eventually released from the membrane, Rab7 is recruited towards the endosome membrane by the Rab7GEF activity of BuMC1. However, recent evidence suggests that BuMC1 is not only required for Rab7 recruitment but also for the inactivation or membrane release of Rab5, though the mechanistic details remain unexplored (Borchers et al., 2023; Herrmann et al., 2023a). Because mutation of any of the BuMC1 proteins causes Rab5 accumulation in endosomes, this raises the question of whether the recruitment of Rab7 is also necessary for an efficient Rab5 release.

Therefore, we investigated the direct role of Rab7 in releasing Rab5 from EEs in nephrocytes (Figure 5) by expressing variants of Rab7. Surprisingly, overexpression of wildtype Rab7 in nephrocytes causes early lethality in our experiments, preventing us from studying its effects on late larval nephrocytes. Thus, we induced expression of constitutively active Rab7 (Rab7^CA^; Q67L), a GTP-locked version of Rab7 that remains at the LE/MVB membrane and fails to be inactivated by GAP-mediated hydrolysis (Bucci et al., 2000). Nephrocytes were stained for Rab5 and Rab7, and we found that the vesicle appearance, size, and the localisation of Rab5 and Rab7 appear similar to that in wildtype cells (Figure 5B).

**Figure 5:**
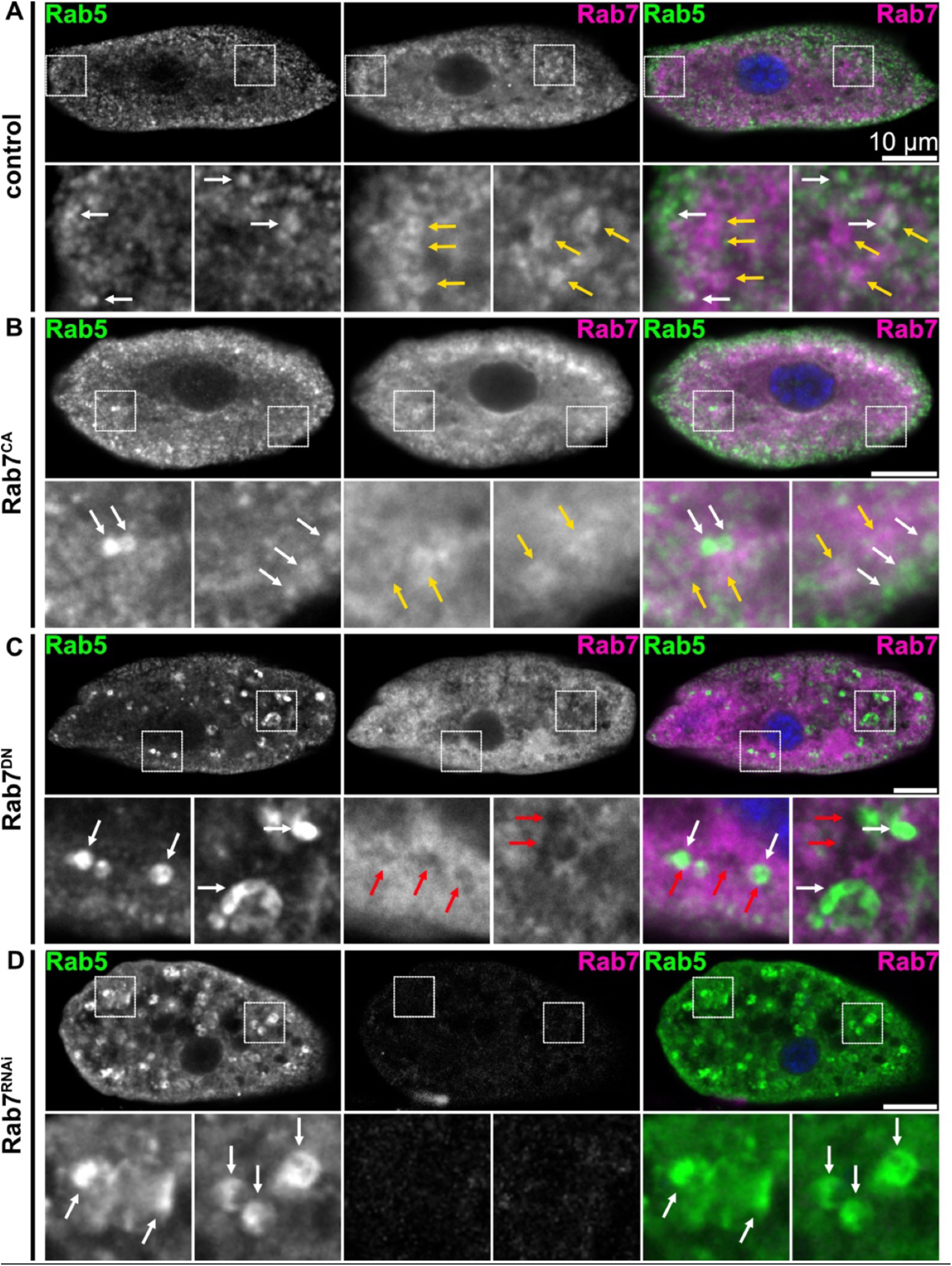
Expression of GDP-locked Rab7, Rab7^DN^, mimics the BuMC1 phenotypes. Nephrocytes from dissected 3^rd^ instar larvae of control flies (A), or larvae of flies in which either Rab7^CA^ (B), Rab7^DN^ (C) or Rab7^RNAi^ (D) was expressed on top of the endogenous Rab5. Nephrocytes were stained with Rab5 and Rab7 specific antibodies. Expression of the three constructs, YFP.Rab7^CA^, YFP.Rab7^DN^ and Rab7^RNAi^ were driven by *hand*C-Gal4. Single channels, merged images and close-ups (indicated by dotted-lined boxes in the overviews) were shown. Enlarged endocytic vesicles were found in flies, in which either Rab7^DN^ was expressed or in which Rab7 was downregulated by RNAi (white arrows). Scale bar indicates 10 μm.

Finally, we examined the effects of overexpressing a GDP-locked dominant-negative Rab7 (Rab7^DN^; T22N) in an otherwise wildtype background. Rab7^DN^ cannot switch from the inactive to the active GTP-bound state. Expression of Rab7^DN^ results in a diffuse endogenous Rab7 signal, indicating that the presence of GDP-locked Rab7 protein impairs the proper localisation of endogenous Rab7 on late endosomal membranes. Importantly, under these conditions, we observed enlarged Rab5 endosomes, with reduced or absent Rab7 decoration (Figure 5, red arrows). These enlarged endosomes may internalise and accumulate Rab5, as we observed in BuMC1 mutants. However, the image resolution is not adequate to ultimately assess this (Figure 5C). Therefore, we assume that the expression of Rab7^DN^ indeed imposes a dominant effect on Rab7 recruitment and thus prevents an efficient switch from Rab5 to Rab7 on the endosome membrane. The inability of Rab7^DN^ to be recruited and activated on endosomal membranes causes the formation of enlarged endosomes that mimic the BuMC1 phenotype. These observations prompted us to test whether the depletion of Rab7 causes similar, characteristic BuMC1 phenotypes in nephrocytes as well. Therefore, we used RNAi-mediated knockdown of Rab7 and stained the nephrocytes for Rab5 and Rab7 (Figure 5D). Based on staining for Rab7, the knockdowns were highly effective, with most of the Rab7 signal absent in Rab7-RNAi-expressing nephrocytes. Importantly, we again observed enlarged vesicles with internalised Rab5 clusters, very reminiscent of BuMC1 mutations or the expression of Rab7^DN^. This finding is further supported by an observation made by Lőrincz and colleagues. In ultrastructural analyses (TEM) of garland nephrocytes in which Rab7 was depleted, they also found enlarged endosomes (Lőrincz et al., 2019).

In summary, our data suggest that effective maturation in nephrocytic endosomes depends on the proper clearance or inactivation of Rab5, as well as on the presence and membrane recruitment of Rab7. Based on the similarities of the observed phenotypes, the BuMC1 complex may be involved in both processes and, therefore, constitutes a major player in the release of Rab5 and recruitment of Rab7. Nevertheless, the in-depth mechanism by which BuMC1 facilitates Rab7 recruitment and activation, as well as Rab5 inactivation, remains unclear.

### Genetic screening for GAPs controlling endosomal maturation in nephrocytes

Based on the observation that cells with a mutation for any of the BuMC1 components fail to inactivate Rab5, we hypothesised that the BuMC1-Rab7 complex might be directly or indirectly involved in the recruitment of an unknown Rab5-GAP to facilitate Rab5 release. Thus, inactivation of such a Rab-GAP should at least partially mimic the BuMC1 phenotype. The *Drosophila* genome harbours 25 potential RabGAPs, classified as GAP proteins based on the presence of a conserved Tre2/Bub2/Cdc16 (TBC) domain (Albert et al., 1999; Fukuda, 2011; Lamber et al., 2019) (Table 1). For some of these potential GAPs, GAP activity has already been shown in *D. melanogaster*, *C. elegans*, yeast, or cultured cells. Known Rab5-GAPs include TBC1D18 (Hiragi et al., 2022), RN-tre (Lanzetti et al., 2000), RUTBC3 (Haas et al., 2005), and *C. elegans* TBC-2 (Chotard et al., 2010; Li et al., 2009). Known Rab7-GAPs include TBC1D5, which has been identified in mammalian cell culture (Jimenez-Orgaz et al., 2018). In the absence of TBC1D5, the Rab7 activity state and localisation are no longer controlled, causing hyperactive Rab7 expanding over the entire endolysosomal system (Jimenez-Orgaz et al., 2018). TBC1D5 has been initially identified as a GAP component of the retromer complex (Roy et al., 2017; Seaman et al., 2009; Zhang et al., 2005). Recently, the yeast GTPase-activating protein Gyp7 (in flies, TBC1D15-17) was shown to regulate Rab7 (Ypt7 in yeast) on late endosomes (Füllbrunn et al., 2024). In the presence of sufficient GDI, Gyp7 activity shifts Ypt7 localisation from the lysosome membrane (the vacuole in yeast) to endosomes (Cabrera and Ungermann, 2013). However, many GAP proteins harbour relatively low substrate specificity, and Gyp7 and TBC1D5 may act on other Rabs in addition to Rab7. Moreover, the depletion of RabGAPs is often compensated by other GAPs, due to the generally promiscuous activity of GAPs against different Rab proteins.

**Table 1:**
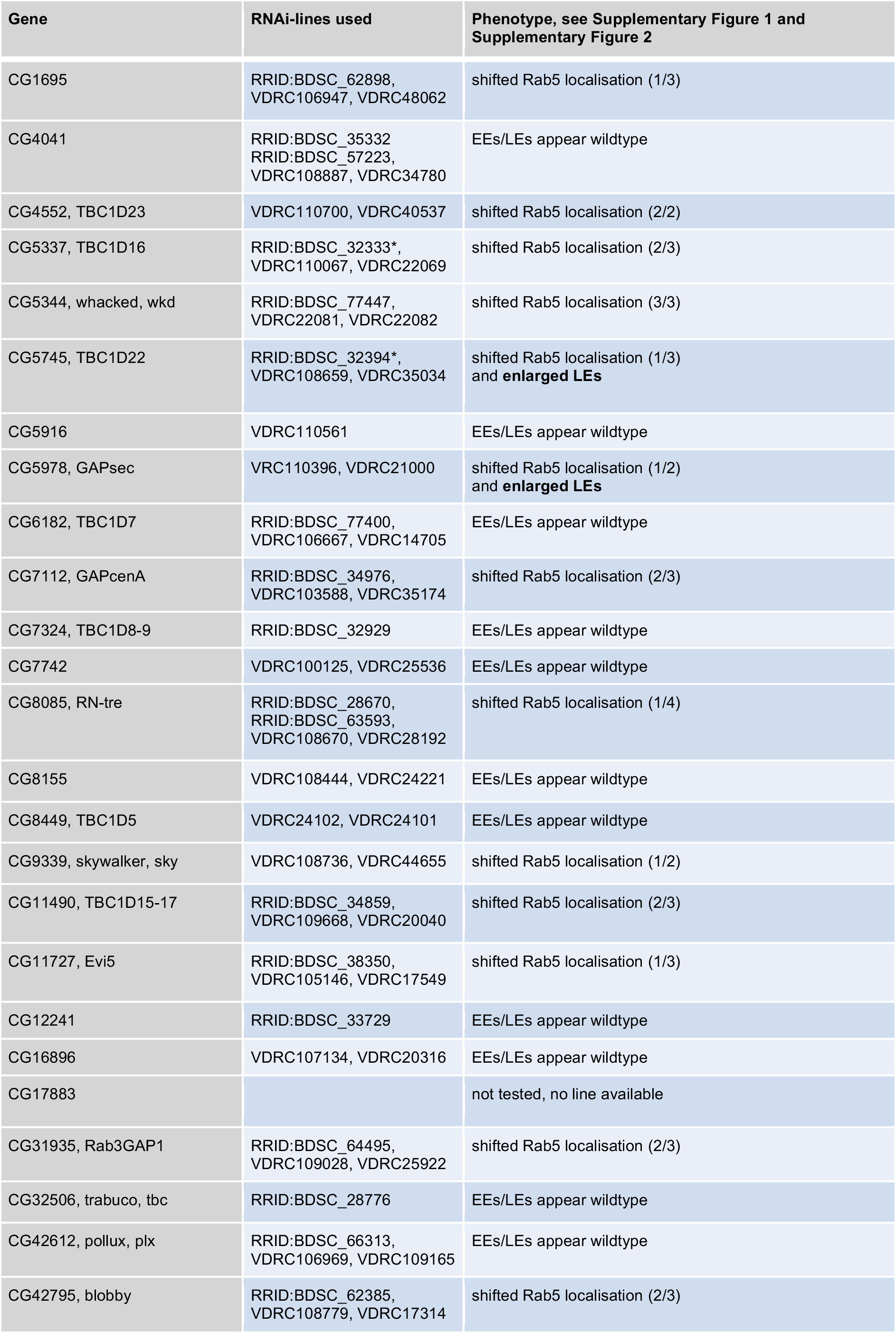
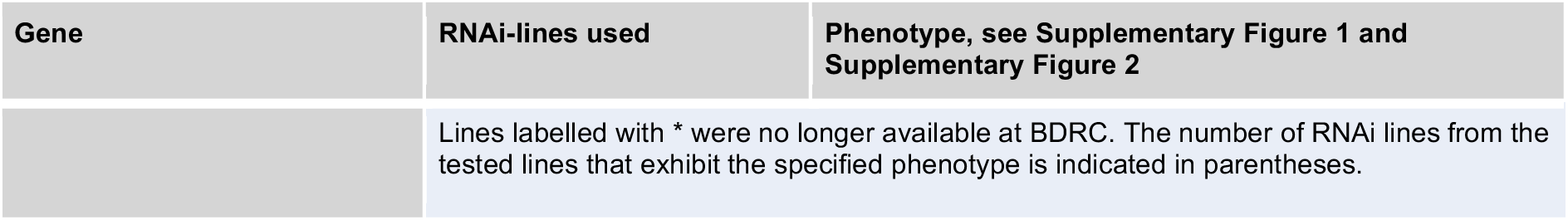
Genetic screening for GAP proteins controlling endosomal maturation in nephrocytes. We tested all annotated potential GAP genes in the *Drosophila* genome for a possible role in endosomal maturation in nephrocytes. One candidate is missing due to the lack of suitable lines for RNAi-mediated downregulation. Pericardial nephrocytes from dissected 3^rd^ instar larvae of control flies and of flies, in which GAPsec or Tbc1d22 was down-regulated by different RNAi-constructs, were stained for Rab5 (anti-Rab5) and YFP.Rab7 (anti-GFP) to identify potential endosomal maturation defects. The RNAi lines used in this study, along with the results of our stainings, are listed. Images of nephrocytes stained for Rab5 and Rab7 are shown in Supplementary Figure S1.

To identify potential GAPs that might be involved in nephrocytic endosome maturation, we performed an unbiased RNAi-mediated knockdown screen (Table 1). Nephrocytes respond well to RNAi-mediated knockdown of endosomal components and, therefore, are highly suitable for a systematic screen. However, one has to take into account here that over 90% of all available RNAi lines exhibit residual activity of 25% or more (Heigwer et al., 2018; Perkins et al., 2015). On the one hand, this is an advantage for studying genes that are essential for the cell. On the other hand, it can be a disadvantage if gene products are only needed in small quantities and therefore a low remaining expression activity is sufficient to sustain gene function. This may lead to weak or absence of phenotypes that would be visible in a complete knockout. We mention this because we observe mild or absent phenotypes with some of the RNAi lines used in our screen.

Expression of RNAi mediating hairpins was controlled by the *hand*-Gal4 driver line, which is active in all nephrocytes at all developmental stages (Paululat and Heinisch, 2012; Sellin et al., 2006). All knockdowns were additionally driven in the presence of TI{TI}Rab7^EYFP^, a functional protein trap in the Rab7 locus (Dunst et al., 2015). To identify endosomal maturation defects, nephrocytes were fixed and stained for Rab5 and YFP.Rab7, and analysed under a cLSM. All RNAi lines available at the time of this study were tested. As an initial read-out, we screened for anomalies in Rab5 and YFP.Rab7 distribution, vesicle size, and the clearance of Rab5 from endosomes. We aimed to identify phenotypes similar to our observations in BuMC1 mutants.

Out of the 58 RNAi lines tested, none resulted in the aberrant accumulation of Rab5 in endosomes (Supplementary Figure 1). Given the nature of RNAi-mediated knockdown screens, with the majority of RNAi lines exhibiting residual 25% or more transcriptional activity (Heigwer et al., 2018), amorphic phenotypes are rarely observed. Moreover, RabGAPs, like several other components of the endocytic pathway, exhibit some functional redundancy. Therefore, a complete loss-of-function phenotype is rather unlikely. However, we succeeded in observing alterations in endosome morphology or Rab5 localisation in eight different potential RabGAP-encoding genes (Table 1). The screen yielded two distinct classes of phenotypes.

In wildtype nephrocytes, Rab5-positive early endosomes predominantly localise directly below the cell cortex. The first phenotypic group, comprising the genes *blobby*, *CG1695*, *Evi5*, *GAPCenA*, *Rab3GAP1*, *RN-tre*, *skywalker*, *TBC1D16*, *TBC1D15-17*, *TBC1D22*, *TBC1D23*, and *whacked*, exhibited a diffuse mis-distribution of Rab5-positive vesicles (Supplementary Figure 1, Supplementary Figure 2, Table 1). To estimate the distribution of Rab5-positive vesicles in cells, we measured the intensity of a Rab5 immunostaining within a 20-pixel-wide and 10 µm-long corridor between the cell cortex and the centre of the cell (Supplementary Figure 2). In wildtype nephrocytes, Rab5 concentration, and thus vesicle density, is highest right below the plasma membrane, and steadily diminishes within the first 5 µm towards the cell centre (Supplementary Figure 2).

The second phenotypic group partially mimicked the BuMC1 phenotype and was observed in cells with knockdown of GAPsec or Tbc1d22 (Supplementary Figure 1). Although no Rab5 clustering was observed, knockdown of either gene resulted in the appearance of unusually large YFP.Rab7-decorated endosomes (Figure 6A-C). Since the appearance of large late endosomes is also seen in BuMC1 mutants, this observation prompted us to investigate both genes in more detail. In *Drosophila*, Tbc1d22 is essential for regulating lipid homeostasis, resulting in lipid droplet accumulation in multiple tissues when it is depleted from cells. Tbc1d22 exhibits GTPase-activating protein (GAP) activity, while the primary substrate in lipid homeostasis is Rab40 (Duan et al., 2021). GAPsec (GTPase activating protein, SECIS-dependent read-through) is the orthologue of the *C. elegans* FAPsec (F45E6.3; NM_077694) (Hirosawa-Takamori et al., 2009) and the human hGAPsec (TBC1D13; PMC3470861).

**Figure 6:**
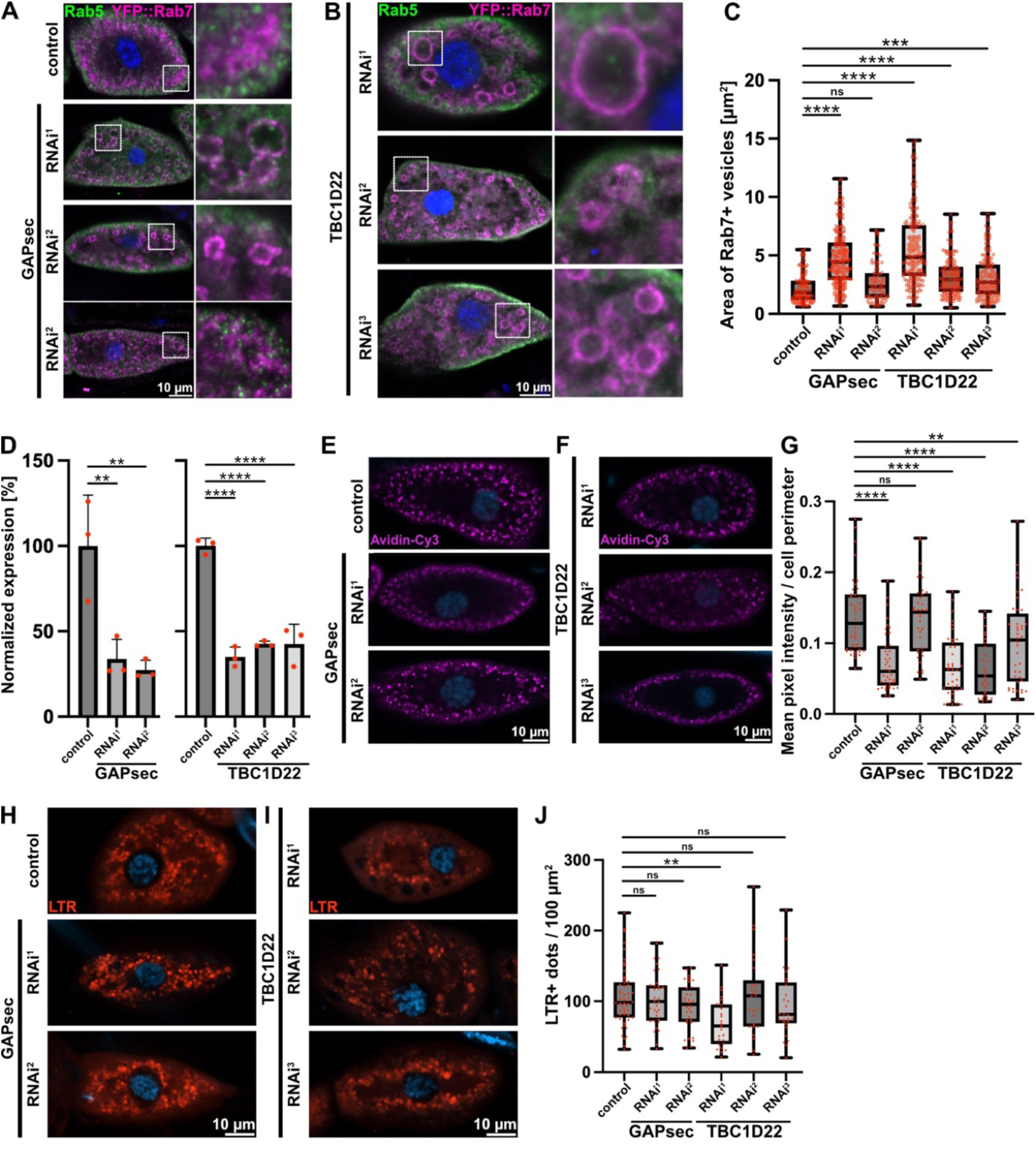
RNAi-mediated downregulation of Tbc1d22 and GAPsec led to the formation of highly enlarged late endosomes. Pericardial nephrocytes from dissected 3rd instar larvae of control flies and of flies in which GAPsec (A) or Tbc1d22 (B) expression was down-regulated by different RNAi constructs and stained for Rab5 (anti-Rab5) and Rab7 (anti-YFP.Rab7) to identify potential endosomal maturation defects. (C) Quantification of Rab7-positive vesicle area. In both cases, highly enlarged Rab7-positive vesicles were detectable. (D) Verification of the RNAi-mediated downregulation capacity of the RNAi lines used. (E) Pericardial nephrocytes from dissected 3^rd^ instar larvae of control flies and of flies, in which GAPsec (E) or Tbc1d22 (F) was down-regulated by different RNAi-constructs, were treated (pulse-chase experiment) with Avidin-Cy3 and imaged after 10 minutes. (G) Quantification of the mean pixel intensity in relation to the cell perimeter of Avidin-Cy3 positive compartments after 10 minutes of incubation with the tracer. (H) Pericardial nephrocytes from dissected 3^rd^ instar larvae of control flies and of flies, in which GAPsec (H) or Tbc1d22 (I) was down-regulated by different RNAi-constructs, were treated with LysoTracker. (J) Quantification of LysoTracker positive dots normalised to a cell area of 100 µm^2^. Mean pixel intensity/perimeter measurements of Lysotracker positive compartments after 10 minutes of incubation with the tracer.

### The role of Tbc1d22 and GAPsec in endosome biogenesis

To determine their possible role in endosome biogenesis, we first assessed the knockdown efficiency of all Tbc1d22 and GAPsec RNAi lines using quantitative PCR (qPCR). Systemic knockdown, driven by *daughterless*-Gal4, resulted in a reduction of the transcripts by about 50-75% (Figure 6D). Therefore, we consider the observed alterations in endocytosis as hypomorphs, rather than a loss-of-function phenotype. Furthermore, although the measured depletion of transcript levels in the entire animal suggests a rather efficient knockdown, we cannot assure that the reduction of gene expression is similar in every single cell or tissue of the animal. As a consequence, the penetrance and severity of the observed phenotypes appeared to be highly variable for certain hairpins.

In the first set of experiments, we measured the size of individual YFP.Rab7-decorated vesicles. Consistent with the initial observation, we found a significant enlargement of late endosomes in GAPsec- and Tbc1d22-knockdown cells. While in control cells, late endosomes rarely reach sizes of more than 5 µm^2^, depletion of either gene can result in vesicles up to 10 - 15 µm^2^ (Figure 6C). Mutations in *bulli* cause a decreased endocytosis rate in nephrocytes, likely due to „backlogging” of cargo (Dehnen et al., 2020). To monitor the rate of endocytosis in GAPsec- and Tbc1d22-knockdown nephrocytes, we performed pulse-chase experiments using Avidin-Cy3 as a tracer molecule with a 1 min pulse and 9 min chase period. We found that the uptake of Avidin-Cy3 is slowed down in both GAPsec and Tbc1d22 knockdowns (Figure 6E-G). This indicates that the GAP depletion results in less efficient endocytic uptake, in agreement with the observed malformation of later endocytic carriers.

We therefore asked next whether the GAP depletions impaired lysosome biogenesis. To specifically label lysosomal compartments in wildtype and mutant nephrocytes, we labelled lysosomes with lysotracker and counted the number of lysosomes in a central focal plane (Figure 6H-J). Except for one Tbc1d22 hairpin, we found that the number of lysosomes was not significantly affected, indicating that the biogenesis of these organelles is not impaired by either GAP depletion (Figure 6J). Taken together, we conclude that both GAPSec and Tbc1d22 function in the maturation of endosomes and therefore regulate endocytosis in nephrocytes, most likely by functioning as a RabGAP.

### Downregulation of GAPsec in nephrocytes causes an increase of abnormally enlarged endosomes

As with the individual components of the BuMC1 complex, we also examined the nephrocytes of animals with downregulated GAPsec using ultrastructural analyses (Figure 7). We found enlarged endosomes, although not as frequently as observed in *mon1*, *ccz1*, or *bulli* mutants (Figure 2). This reduced severity may be due to residual GAPsec activity or to redundancies, i.e., the activity of other GAPs. The labyrinth channel system and the slit diaphragms of the nephrocytes appear normal in our ultrastructural analyses.

**Figure 7:**
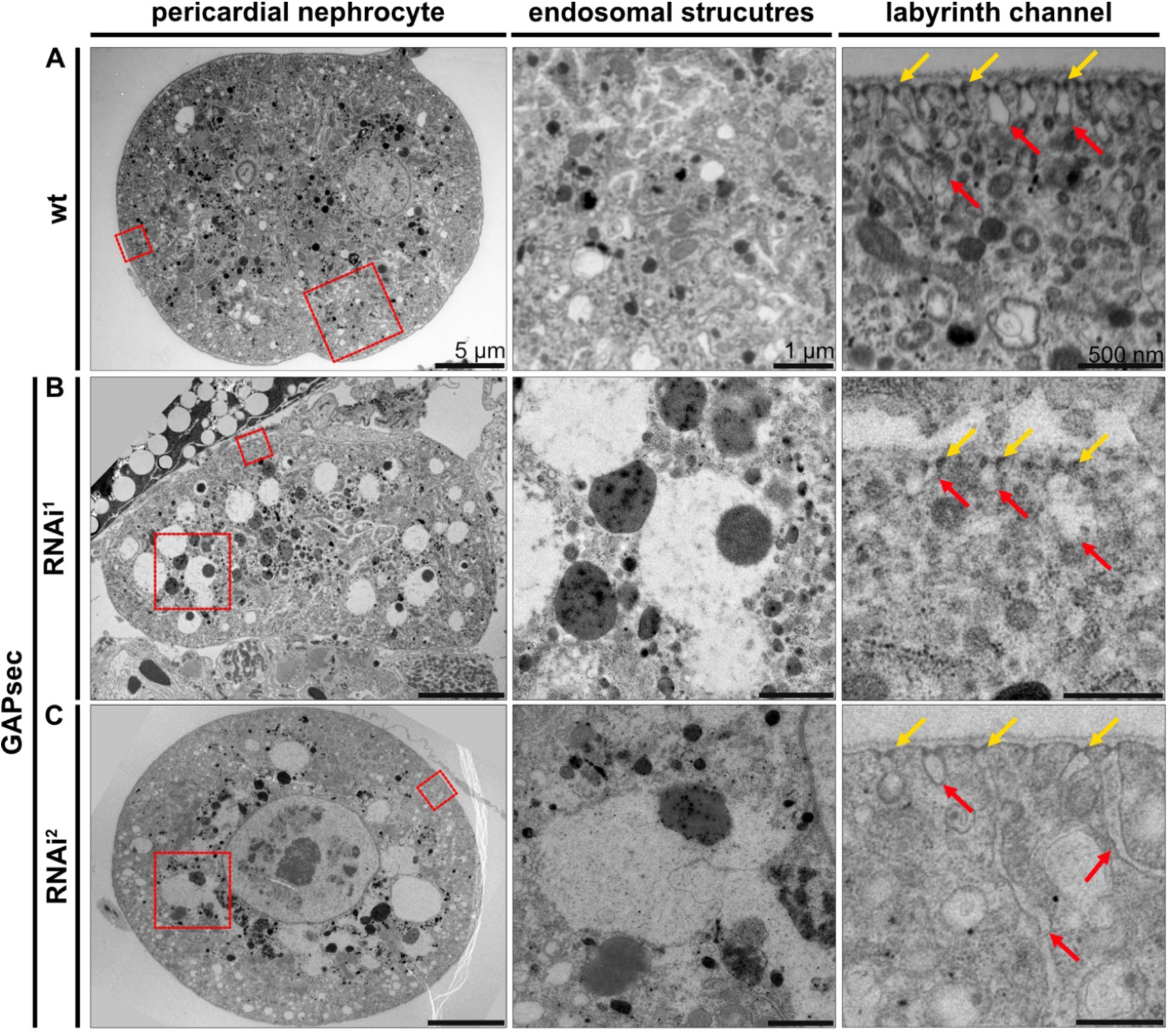
Ultrastructural defects in GAPsec depleted nephrocytes. Ultrastructural analyses of nephrocytes from dissected 3^rd^ instar larvae of control flies (A) and flies, in which GAPsec was down-regulated by two individual RNAi-lines (B, C). Enlarged endosomes are observed in the knock-down cells but not in control nephrocytes. The slit diaphragms (yellow arrows) and the labyrinth channel system (red arrows) are largely unaffected.

### GAPsec has GAP activity for Rab5

Finally, we tested whether GAPsec or Tbc1d22 had GAP activity for either Rab5 or Rab7. Tbc1d22 proved to be unstable even at low concentrations after expression in bacteria, and we thus failed to test for any GAP activity in our assay. Therefore, we focused on GAPSec. We recombinantly expressed Rab5, Rab7, and GAPSec in *E. coli,* and isolated the proteins via affinity purification and size exclusion chromatography. Enrichment and purification for GAPSec is shown in Figure 8A. GTP hydrolysis was then measured via an HPLC-based multi-turnover assay (Figure 8B). Rab5 and Rab7 exhibited comparable low intrinsic GTP hydrolysis rates. Upon addition of GAPsec, Rab5, but not Rab7, hydrolysed GTP efficiently, suggesting that GAPsec is a new Rab5GAP in *Drosophila*.

**Figure 8:**
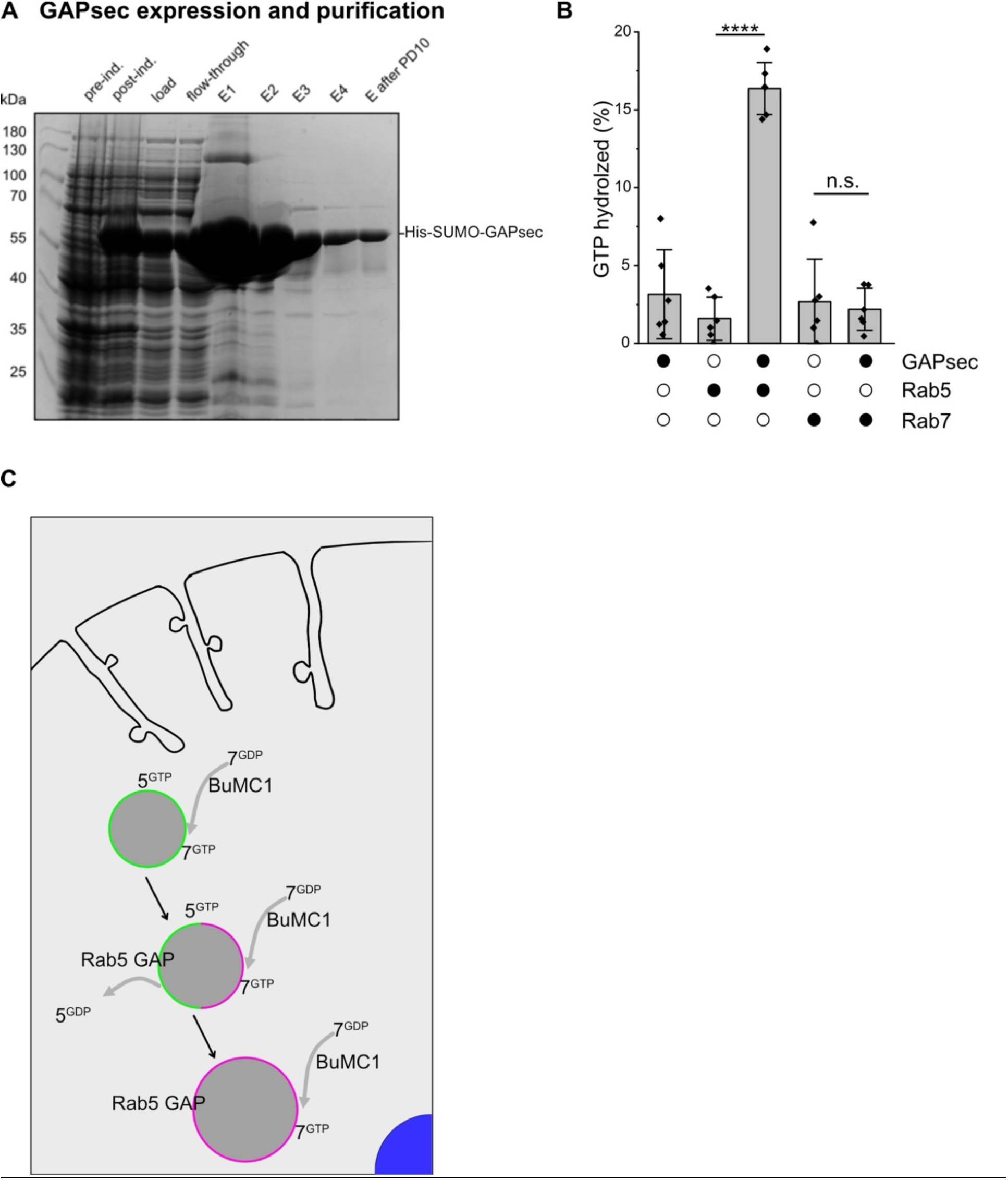
GAPsec activity converts Rab5GTP to Rab5GDP, thus it acts as a Rab5 GAP in Drosophila. (A) SDS-PAGE gel of purified proteins used for *in vitro* assays. (B) GTP hydrolysis assay of Rab5 and Rab7 in the absence and presence of GAPsec. Data are presented as means ± standard deviation from n=6 independent repeats. Statistical analysis by two-sided t-test; **** p<0.0001, n.s. not significant. (B) shows a possible mode of action for BuMC1 and GAPsec activity in nephrocyteś endosomal pathway. Active Rab5 on the membrane recruits and activates BuMC1, which in turn promotes Rab7 activation. BuMC1 also triggers the activity of the Rab5 GAP (GAPsec), leading to Rab5 inactivation and its release from the membrane. Rab5^GTP^ on the endosomal membrane is indicated in green, and Rab7^GTP^ is labelled in magenta.

Taken together, our data reveal GAPsec as a new Rab5GAP protein involved in endocytosis in *Drosophila* nephrocytes.

## DISCUSSION

Our data suggest that the activity of the trimeric BuMC1 complex highly regulates the early-to-late transition of endosomal maturation in several ways. On the one hand, it acts as the major Rab7-GEF that recruits Rab7 towards endosomal membranes and initiates MVB/LE maturation into lysosomes (Borchers et al., 2023; Dehnen et al., 2020; Herrmann et al., 2023a; Herrmann et al., 2023b; Langemeyer et al., 2020). The structure of the metazoan Rab7-GEF complex Bulli-Mon1-Ccz1 (BuMC1) (Herrmann et al., 2023b; Yong et al., 2023) shows that the metazoan-specific subunit Bulli interacts directly with Ccz1 but not with Mon1. When Bulli is deleted, the *in vitro* GEF activity of the dimeric complex is slightly reduced, which can be decisive in a physiological context, where localisation to endosomes depends on multiple membrane interactions (Borchers et al., 2023; Dehnen et al., 2020; Langemeyer et al., 2020; Wilmes et al., 2025).

On the other hand, BuMC1 directly interacts with Rab5 at early endosomes via a Mon1 longin domain (Borchers et al., 2023; Herrmann et al., 2023b), indicating a potential role of the metazoan BuMC1 complex at earlier stages of endosomal maturation. The unstructured N-terminal part of Mon1 likely folds back to the core of Mon1, resulting in autoinhibition of the BuMC1 GEF activity (Borchers et al., 2023). When Rab5 binds to a conserved site in Mon1, the BuMC1 complex is both recruited to membranes and activated, driving the nucleotide exchange of Rab7 (Borchers et al., 2023). Several open questions remain. When, where, and by which GAP is Rab5 inactivated? How is the Rab5 GAP recruited, and how is this process regulated to prevent premature Rab5 to Rab7 turnover?

A key observation from our analysis of the *bulli*, *mon1*, and *ccz1* mutants is defective endosomal maturation, which is characterised by the formation of giant endosomes. These endosomes accumulate Rab5 internally and fail to become Rab7 positive. Thus, they cannot acidify, and maturation into a functional lysosome is impaired. We have observed that the phenotypes of the individual BuMC1 mutants and other effectors of the endosomal maturation pathway are not identical, but sometimes merely similar. This is not surprising and can be explained in different ways. On the one hand all players may act in a number of cellular pathways, beyond EE-LE maturation. It is therefore conceivable that the knockout phenotypes vary to some extent. However, they remain consistent in some essential aspects, including the presence of enlarged endosomes. On the other hand, it was shown that Bulli is important for membrane binding (Wilmes et al., 2025), whereas Mon1 and Ccz1 are necessary for the GEF activity (Dehnen et al., 2020; Langemeyer et al., 2020). Thus, loss of Bulli results in disrupted membrane association, while absence of Mon1 or Ccz1 lead to impaired activity of the BuMC1 complex, likely explaining the difference in phenotypic manifestation. Furthermore, the mutations are of different genetic nature. *Mon1*^mut4^ is a premature stop (Q70term) amorphic allele (Yousefian et al., 2013), *ccz1*^d113^ is a 1644bp deletion that removes almost all of the *ccz1* coding region (Hegedűs et al., 2016), and *bulli*^6-61^ is the result of a CRISPR-Cas9 mediated 8 bp deletion within the coding region causing a frame shift and a premature stop codon (Dehnen et al., 2020). We therefore propose that the individual genetic characteristics of the various mutations may contribute to variations in the observed phenotypes.

However, the BuMC1 phenotype can be reproduced or mimicked by manipulating Rab5 and Rab7 (as shown in this work). First, when a GTP-locked version of Rab5, Rab5^CA,^ is expressed in nephrocytes, endogenous Rab5 becomes internalised upon endosomal maturation (Figure 3B). Moreover, expression of GDP-locked Rab7, Rab7^DN^, and depletion of Rab7 (via Rab7^RNAi^) inhibit the endosomal maturation pathway (Figure 4C, D). Mutant cells have enlarged endosomes, where they accumulate Rab5 in intraluminal vesicles. This shows that the regulation of the activation and inactivation of Rab proteins, such as Rab5 and Rab7, is impaired. The Rab7 GEF activity of BuMC1 is thus essential for the progression of endosomal maturation. In addition, BuMC1 interacts directly with Rab5 at early endosomes and may be involved in the inactivation and release of Rab5 (Borchers et al., 2023; Herrmann et al., 2023a; Herrmann et al., 2023b).

Furthermore, our results indicate that, in the absence of Rab7, vesicle fusion is misregulated, resulting in enlarged vesicles as mentioned above. These vesicles are unlikely to mature into lysosomes. The expression of a constitutively active Rab7^CA^ is tolerated by nephrocytes, which is likely due to the remaining endogenous wildtype Rab7. In contrast, expression of a dominant-negative form of Rab7^DN^ resulted in enlarged vesicles with internalised Rab5. We speculate that Rab7^DN^ cannot be recruited to endosomes, but blocks the recruiter (BuMC1), thus mimicking an absence of Rab7 (Figure 5C). As a result, RNAi-mediated downregulation of *Rab7* causes endosomal defects (Figure 5D), similar to those found in *bulli*, *mon1,* and *ccz1* mutants, and when Rab5^CA^ is overexpressed (Figure 1, Figure 4B).

Constitutively active Rab5^CA^ stays at the EE membrane and inhibits endosomal maturation. Rab5-GTP needs to be inactivated for proper membrane release. Cells possess GAPs for this task. More than 20 GAPs are encoded in the *Drosophila* genome, but the substrate and function of only a few of them are known. Our RNAi-based screen identified two Rab5 GAP candidates, Tbc1d22 and GAPsec. Depletion of either protein led to greatly enlarged endosomes (Figure 6, Table 1), similar to BuMC1 mutants, or Rab5^CA^, and Rab7^DN^ expressing nephrocytes (Figure 1, Figure 2, Figure 4, Figure 5), and GAPsec has indeed Rab5-specific GAP activity (Figure 8). Because the GAPsec knockdown showed a distinct phenotype compared to loss of BuMC1 subunits regarding the intraluminal accumulation of Rab5, it is unlikely that it is the only Rab5-GAP in *Drosophila* nephrocytes required for endosomal maturation. However, none of the previously reported Rab5-GAPs showed any effect in our siRNA screen. Therefore, multiple GAPs might act redundantly to inactivate Rab5. Whether BuMC1 or Rab7 are directly involved in the recruitment of GAPsec or other Rab5-GAPs to endosomal membranes needs to be tested in future studies.

Taken together, the results show that the transition from Rab5 to Rab7 on endosomes, and thus endosomal maturation, is regulated by BuMC1. Our model postulates that BuMC1 is required for Rab7 activation, which in turn enables the proper function of a Rab5 GAP (GAPsec) and thereby promotes the inactivation of Rab5^GTP^ (Figure 8C).

## MATERIALS AND METHODS

### Fly stocks and genetics

The following fly stocks were used in this study: *bulli*^6-61^ (Dehnen et al., 2020) and *hand*C-Gal4 (Sellin et al., 2006) from our laboratory, *mon1*^mut4^ was a gift from Thomas Klein, Düsseldorf, Germany (Yousefian et al., 2013), *ccz1*^d113^ was a gift from Gábor Juhász, Szeged, Hungary (Hegedűs et al., 2016). Lines obtained from the Bloomington Drosophila Stock Center (BDSC) and Vienna Drosophila Resource Center (VDRC): control flies w^1118^ (RRID:BDSC_3605), UAS-YFP.Rab5^WT^ (RRID:BDSC_24616), UAS-YFP.Rab5^CA^ (RRID:BDSC_9773), UAS-YFP.Rab5^DN^ (RRID:BDSC_9771), UAS-YFP.Rab7^CA^ (RRID:BDSC_24103), UAS-YFP.Rab7^DN^ (RRID:BDSC_9778), UAS-Rab7^RNAi^ (RRID:BDSC_27051), eYFP.Rab7 (RRID:BDSC_62545) (we used this line for our GAP-Screen), UAS-GAPsec-RNAi^1^ (VDRC_21000), UAS-GAPsec-RNAi^2^ (VDRC_110396), UAS-Tbc1d22-RNAi^1^ (RRID:BDSC_32394) (this line was not longer maintained at Bloomington), UAS-Tbc1d22-RNAi^2^ (VDRC_35034), UAS-Tbc1d22-RNAi^3^ (VDRC_108659), UAS-YFP.Rab5^CA^ (RRID:BDSC_9774), *da*-Gal4 (RRID:BDSC_95282).

### GAP screen, fly stocks used

See Figure 5, a minimum of 10 cells from three larvae were imaged.

### Immunohistochemistry

3^rd^ instar larvae were dissected in PBS and fixed with 4 % paraformaldehyde (PFA) in PBS for one hour at RT. After three washing steps of 10 min, specimens were permeabilised with 1 % Triton X-100 in PBS for one hour at RT, followed by three further washing steps with BBT (0.1 % BSA and 0.1 % Tween-20 in PBS) for 10 min each. Subsequently, specimens were incubated for 60 min in a blocking solution containing 1 % BSA and 0.1 % Tween-20 in PBS followed by incubation with the primary antibodies (rabbit anti-Rab5, 1:250, Abcam Cat# ab31261, RRID:AB_882240, Cambridge, United Kingdom; mouse anti-Rab7, 1:10, Developmental Studies Hybridoma Bank Cat# Rab7, RRID:AB_2722471, University of Iowa, IA, USA; chicken anti-GFP, 1:500, Abcam Cat# ab13970, RRID:AB_300798, Cambridge, United Kingdom; mouse anti-Spectrin, 1:20, Developmental Studies Hybridoma Bank Cat# 3A9, RRID:AB_528473, University of Iowa, IA, USA; mouse anti-Pyd, 1:50, Developmental Studies Hybridoma Bank Cat# PYD2, RRID:AB_2618043, University of Iowa, IA, USA) in BBT overnight at 8 °C. Samples were rinsed three times with BBT (10 min, RT) and incubated with secondary antibodies (anti-rabbit Alexa Fluor 488, 1:200, Jackson ImmunoResearch Laboratories, Inc, West Grove, PA, U.S.A, Code: 115-165-003, RRID:AB_2338680; anti-mouse Cy3, 1:200, Jackson ImmunoResearch Laboratories, Inc, West Grove, PA, U.S.A, Code: 115-165-003, RRID: AB_2338680; anti-chicken Alexa Fluor 488, 1:200, Jackson ImmunoResearch Laboratories, Inc, West Grove, PA, U.S.A, Code: 703-545-155, RRID:AB_2340375) in BBT for two hours at RT followed by three washing steps with BBT. Samples were embedded in Fluoromount-G mounting medium containing DAPI (Thermo Fisher, Waltham, MA, USA). Confocal images were captured with a laser scanning microscope (LSM800, Zeiss, Jena, Germany) equipped with a Zeiss EC Plan-Neofluar 40x / NA 1.30 Oil DIC M27 40x objective, Multikali PMT detector and Zen2.6 software. Filters for Alexa Fluor 488, Cy3 and DAPI were used. In case of Pyd staining, a Zeiss LSM 880 with fast Airyscan equipped with a C Plan-Apochromat 63x / NA 1.4, DIC, oil immersion was used.

### Image processing and analyses

Image processing was done with Fiji (RRID:SCR_002285) (Rueden et al., 2017; Schindelin et al., 2012) and Affinity Photo (Serif, Nottingham, United Kingdom, RRID:SCR_016952). For quantification of Rab7+ vesicles, a minimum of 15 cells from five animals were analysed. An ordinary one-way ANOVA was performed for statistical analysis using GraphPad Prism 9 (RRID:SCR_002798). For quantification of Rab7+ vesicles, a minimum of 15 cells from five animals were analysed. An ordinary one-way ANOVA was performed for statistical analysis. Analysis of Rab5 distribution was performed by generating an intensity profile (10 µm line length; line width: 20 pixels). Measurements were initiated at the outermost cell periphery. The fluorescence intensity values were normalised using the following equation: (x-x_min_) / (x_max_ - x_min_). Normalised intensity values were plotted against the distance of 10 µm. The Kruskal-Wallis test was performed for statistical analysis. The depth of the labyrinth channels was quantified based on the spectrin staining. Single slice images of the middle of the cells were captured, and the depth of the labyrinth channels was measured using the straight-line tool implemented in Fiji, from the outside to the inside of the cell. Six different parts of each cell were analysed. At least five cells from a minimum of three animals (3^rd^ instar larvae) were quantified. One-way ANOVA was performed for statistical analyses. Additionally, the relative cell size of wildtype, *bulli*^6-61^, *mon1*^mut4^, and *ccz1*^d113^ pericardial nephrocytes was quantified by measuring the maximum cell area using the polygon selection tool in Fiji. The percentage of the cell area was calculated by setting the median wildtype cell area to 100 % and adjusting all other values accordingly. For statistical analyses, one-way ANOVA was performed. At least 13 cells from a minimum of 9 animals (3^rd^ instar larvae) were quantified.

### Avidin-Cy3 Uptake assay

3^rd^ instar larvae were dissected in artificial haemolymph (Meyer et al., 2024; Vogler and Ocorr, 2009). The preparation buffer was replaced with artificial haemolymph containing 0.02 mg/ml Avidin-Cy3 (#E4142, Sigma-Aldrich, St. Louis, Missouri, USA). Samples were incubated in the staining solution for 1 min (pulse), followed by a 9 min chase period with artificial haemolymph at room temperature in the dark. Specimens were washed with artificial haemolymph and uptake was stopped by fixation with 4% methanol-free formaldehyde in artificial haemolymph. After two brief washing steps with artificial haemolymph, the specimens were embedded in Fluoromount-G mounting medium containing DAPI, and images of pericardial nephrocytes were captured using identical settings (LSM800 (Zeiss, Jena, Germany)). The mean pixel-intensity measurement function provided by the Fiji software package was used to quantify uptake efficiency. Pixel intensity was measured in relation to the perimeter of the cell. At least 28 cells from 10 animals were analysed. For statistical analyses, one-way ANOVA was performed (GraphPad Prism 9, RRID:SCR_002798).

### LysoTracker Red staining

After dissection of the animal, the artificial hemolymph was removed and replaced by LysoTracker solution (0.5 µM LysoTracker™ Red DND-99 (Thermo Scientific™, Waltham, MA, USA) in artificial hemolymph). Specimens were incubated for 15 minutes at room temperature in the dark under moderate shaking, followed by three washing steps with artificial hemolymph (each step lasting 1 minute). Animals were then embedded in Fluoromount-G mounting medium containing DAPI (#00-4959-52, Thermo Scientific™, Waltham, MA, USA) and single-slice images were immediately acquired using an LSM800 (Zeiss, Jena, Germany) with identical imaging settings. Images analysis was performed using Fiji (RRID:SCR_002285). Background subtraction was applied using the “Subtract Background” tool (rolling ball radius: 20 pixels; sliding paraboloid; smoothing disabled). LysoTracker Red positive vesicles were quantified and normalised to the cell area of 100 µm^2^. One-way ANOVA was performed for statistical analyses (GraphPad Prism 9, RRID:SCR_002798). At least 25 cells from 10 animals were analysed.

### Verification of GAPsec and Tbc1d22 RNAi lines

GAPsec and Tbc1d22 RNAi lines (see Figure 6) were tested for efficiency by qRT-PCR. The ubiquitously active *da*-Gal4 driver was used to induce RNAi hairpin expression. Total RNA (1 µg), isolated from 3^rd^ instar larvae (at least eight animals per replicate) using the Direct-zol^TM^ DNA/ RNA MiniPrep Kit (#R2080, Zymo Research, RRID:SCR_008968), was treated with DNase I (Thermo Scientific™, Waltham, MA, USA) according to the manufacturer’s instructions and used as template for cDNA synthesis (Luna Script Reverse Transcriptase, New England Biolabs, Ipswich, MA, USA). qRT-PCR was performed according to standard protocols using the GoTaq® 2-Step RT-qPCR Kit (Promega, Madison, WI, USA) and an iCycler iQ Real-Time PCR System (Bio-Rad, Munich, Germany). Primer pairs were designed using QuantPrime, applying the preset to select regions containing at least one intron and allowing splice variant hits. Data were evaluated as previously described (Simon, 2003). The rp49 gene served as reference. The following primer pairs were used: 5’-CACAAATGGCGCAAGCCCAAG-3’ (rp49 forward),5’-CATTTTTTAACTAAAAGTCCG-3’ (rp49 reverse), 5’- GATCTGATACAGCGCATTGACGTG-3’ (Tbc1d22 forward) and 5’-CTCGCGTGTCAGCAGATTGTTC-3’ (Tbc1d22 reverse), 5’-GTACTGCAAGATCTCAGCATAGCC-3’ (GAPsec forward), 5’-TAGCCAAGGAGTAGCTTCCAACTG-3’ (GAPsec reverse). At least three biological replicates were performed. For statistical analyses, one-way ANOVA was performed using GraphPad Prism 9.

### Transmission electron microscopy

Briefly, specimens were prepared in PBS and subsequently fixed for 4 h at RT in fixative (2% glutaraldehyde (Sigma, Germany) / 4% paraformaldehyde (Merck, Germany) in 0.05 M cacodylate buffer pH 7.4). Next, specimens were post-fixed for 2 h at room temperature in 1% osmium tetroxide in 0.05 M cacodylate buffer pH 7.4 (Sciences Services, Germany) and dehydrated stepwise in a graded ethanol series followed by 100% acetone. Subsequently, specimens were embedded in Epon 812 and polymerised for 48 hours at 60 °C. Ultrathin sections (70 nm) were cut on an ultramicrotome (UC6 and UC7 Leica, Wetzlar, Germany) and mounted on formvar-coated copper slot grids. Sections were stained for 30 minutes in 2% uranyl acetate (Sciences Services, Germany) and 20 minutes in 3 % lead citrate (Roth, Germany). A detailed protocol for processing nephrocytes for transmission electron microscopy (TEM) analysis is available elsewhere (Psathaki et al., 2018). All samples were analysed at 80 kV using a Zeiss 902 and a Zeiss LEO912 transmission electron microscope, and at 200 kV using a Jeol JEM2100-Plus transmission electron microscope (Zeiss, Jena, Germany; Jeol, Tokyo, Japan). TEM preparation of fly tissue was previously described in detail (Psathaki and Paululat, 2022). To calculate the number of slit diaphragms (SDs) in wildtype and in *bulli*, *mon1* and *ccz1* mutant nephrocytes, at least three different areas of the cell were analysed to determine the number of SDs per 5 µm of the cell perimeter. Three cells from three individual animals per genotype were analysed and one-way ANOVA was performed for statistical analyses (GraphPad Prism 9).

### Expression and purification of recombinant proteins

*Drosophila* Rab5, Rab7, and GAPsec were expressed and purified essentially as described (Fullbrunn et al., 2024) with some modifications. In brief, GST-TEV*-*Rab5, GST-TEV-Rab7, and His6-SUMO-GAPsec were expressed in *Escherichia coli* BL21 DE3 (Rosetta) cells. Cells were grown in the presence of the corresponding antibiotics at 37 °C in Luria Broth medium until an OD600 = 0.6, before the addition of 0.5 mM isopropyl-β-d-thiogalactoside induced protein expression. After 16–18 hours of protein expression at 16°C, cells were harvested by centrifugation at 4,000 g, 4°C for 10 minutes. Cells were resuspended in a buffer containing 20 mM Na_2_HPO_4_/NaH_2_PO_4_, pH 7.4, 500 mM NaCl, 5% glycerol, 1 mM DTT, and 1 mM MgCl_2_. During lysis, buffers were supplemented with protease inhibitor mix HP (Serva), 0.025 mg/ml DNase I and 1 mg/ml lysozyme. Cell lysis was performed using a Microfluidiser (Microfluidics, Inc.), and the cell lysate was cleared by centrifugation at 40,000 × *g*, 4°C, for 30 min. The cleared lysate was incubated with nickel-nitriloacetic acid agarose (Qiagen) for purification of His6-SUMO-GAPsec or with glutathione sepharose fast-flow beads (GE Healthcare) for GST-fusion proteins (GST-TEV-Rab5, GST-TEV-Rab7). After incubation for 2 h, 4 °C on a turning wheel, and extensive washing of the beads, His6-SUMO fusion proteins were eluted from the beads with the respective buffers containing 300 mM imidazole and cleaved using Sumo protease for overnight at 4°C (coldroom). GST fusion proteins were cleaved from the beads during incubation with TEV protease for overnight at 4 °C on a turning wheel with one buffer exchange. Samples were further purified with an ENrich^TM^ SEC650 (Bio-rad) size exclusion column equilibrated with 50 mM HEPES, pH 7.4, 150 mM NaCl, 1.5 mM MgCl_2_, and 1 mM DTT. Proteins were snap frozen in liquid nitrogen and stored at -70°C until further use.

### GTP hydrolysis assay

Intrinsic and GAP-stimulated GTPase activity were measured with an HPLC-based assay. 5 µM Rab5 or Rab7 were incubated with and without 15 µM GAP protein (GAPsec) in assay buffer (50 mM HEPES, 150 mM NaCl, pH 7.5) supplemented with 20 mM EDTA and 1 mM DTT. To start the reaction 5 mM MgCl_2_ and 50 µM GTP were added. The samples were briefly mixed, and aliquots were snap-frozen in liquid nitrogen at time points 0 min and 120 min and stored at –70 °C. The assay samples were boiled for 5 min at 95 °C, and the precipitated protein was removed by centrifugation (15 min, 4 °C, 20,000 × *g*). Of the cleared supernatant, 10 µl was injected in an Agilent 1260 Infinity II HPLC system equipped with an autosampler and a DAD HS detector. Analytes were separated on an AMAZE HA mixed phase column (Helix Chromatography). The separation of G-nucleotides was achieved by a stepwise double gradient with increasing buffer (200 mM KH_2_PO_4_, pH 2.0, 30-80 %) and acetonitrile (15-20 %) concentrations. The UV traces at 254 nm were used to monitor the elution of nucleotide. UV traces were semi-automatically analysed with OriginPro 2024 (OriginLab Corporation). Traces were baseline corrected, the peaks corresponding to GDP and GTP integrated and the ratio of hydrolysed GTP was calculated. Individual biological repeats were calculated from three technical repeats.

## Acknowledgements

We thank Martina Biedermann, Mechtild Krabusch, and Kerstin Etzold for excellent technical assistance. This work was supported by grants from the Deutsche Forschungsgemeinschaft (DFG) to Achim Paululat, Daniel Kümmel and Christian Ungermann (PA517/12-2, KU2531/7-1, UN111/9-2), and by the SFB 944 and SFB 1557 (to A. Paululat, D. Kümmel, and C. Ungermann). We also acknowledge the support of the Open Access Publishing Fund of Osnabrück University. We are also grateful to the Vienna Drosophila Resource Centre and the Bloomington Drosophila Stock Centre for providing stocks essential for this work.

## Data and resource availability

All relevant data can be found within the article and its supplementary information.

## Supplementary Figure and File Legends

**Supplementary Figure 1.**
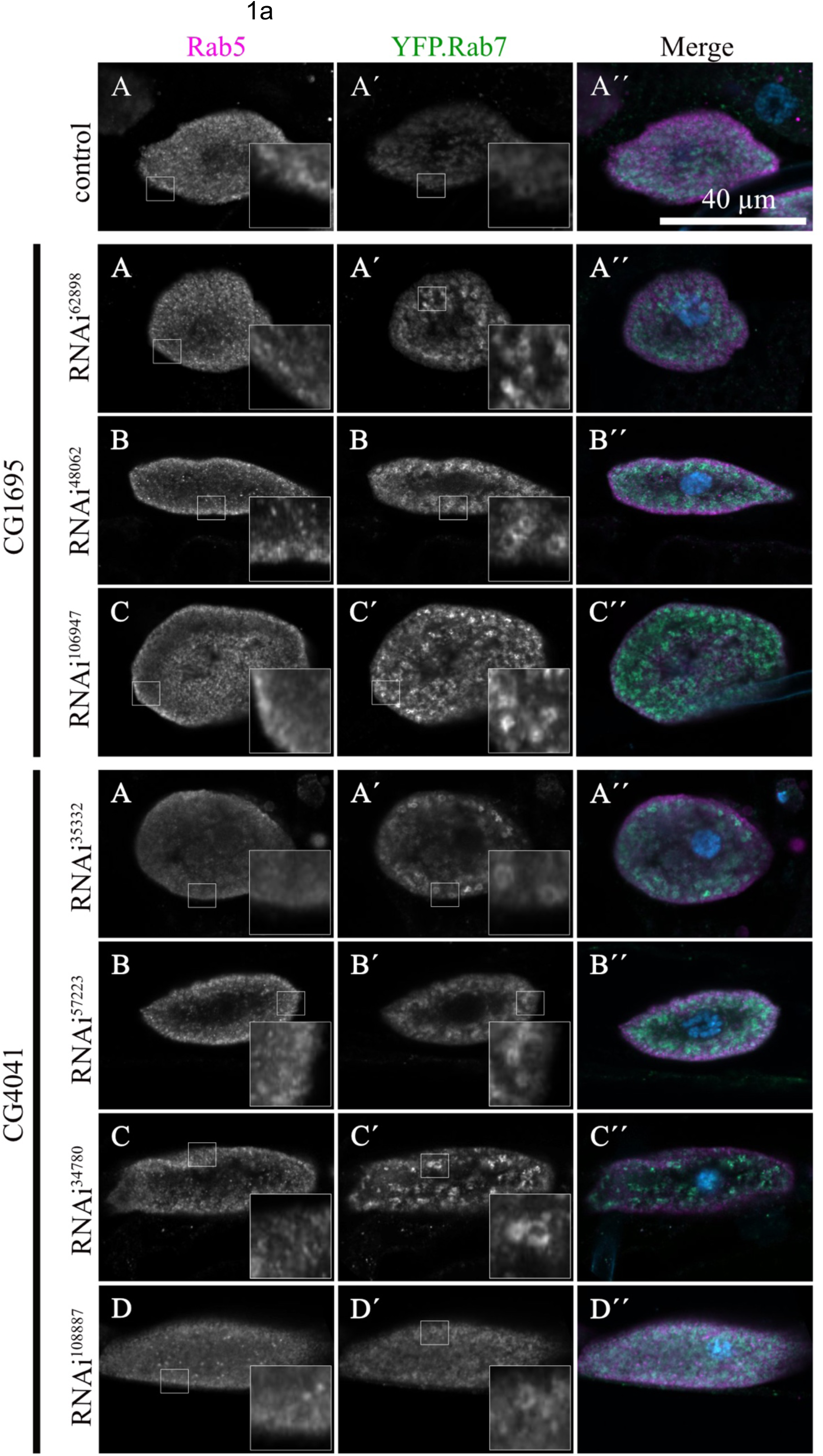
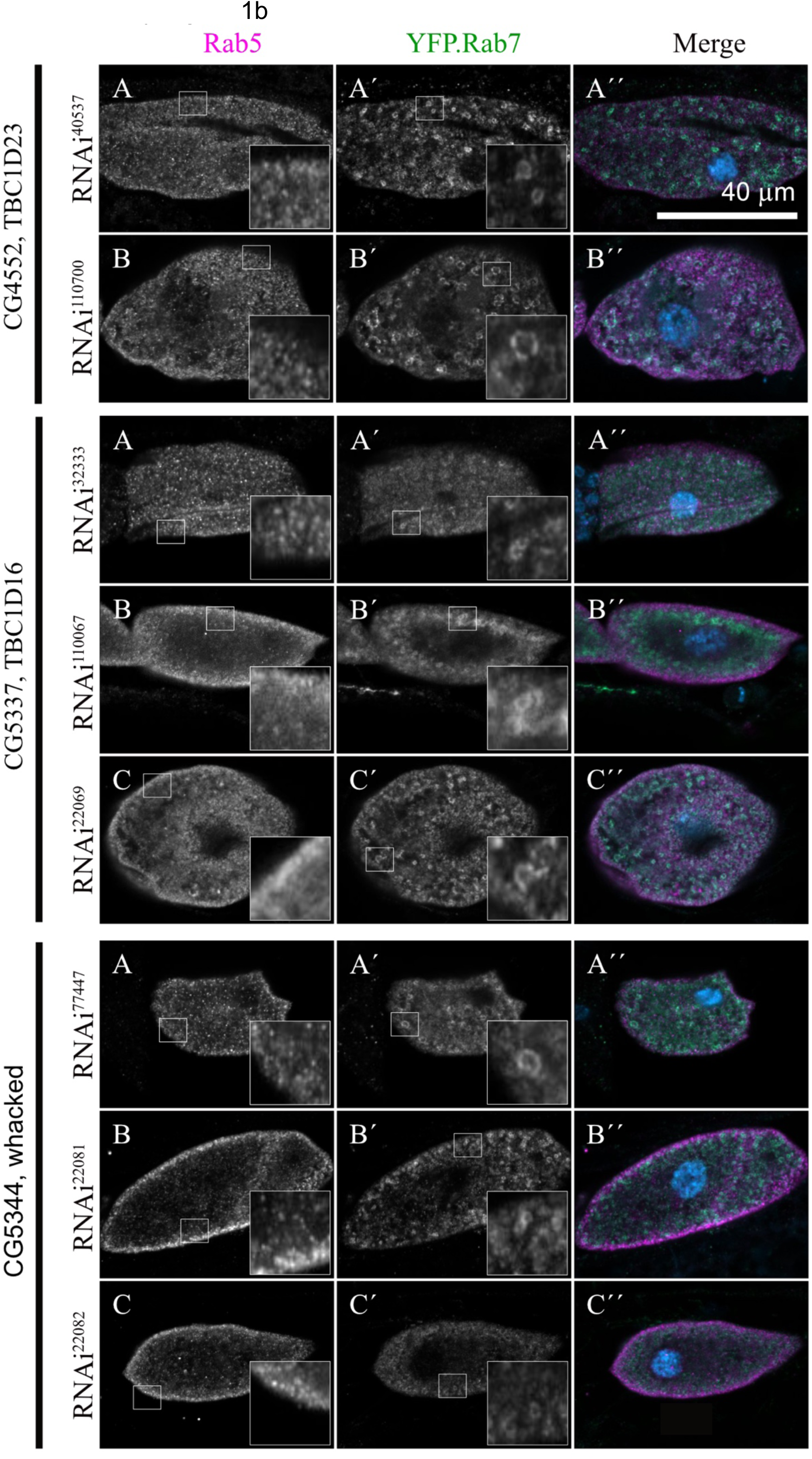
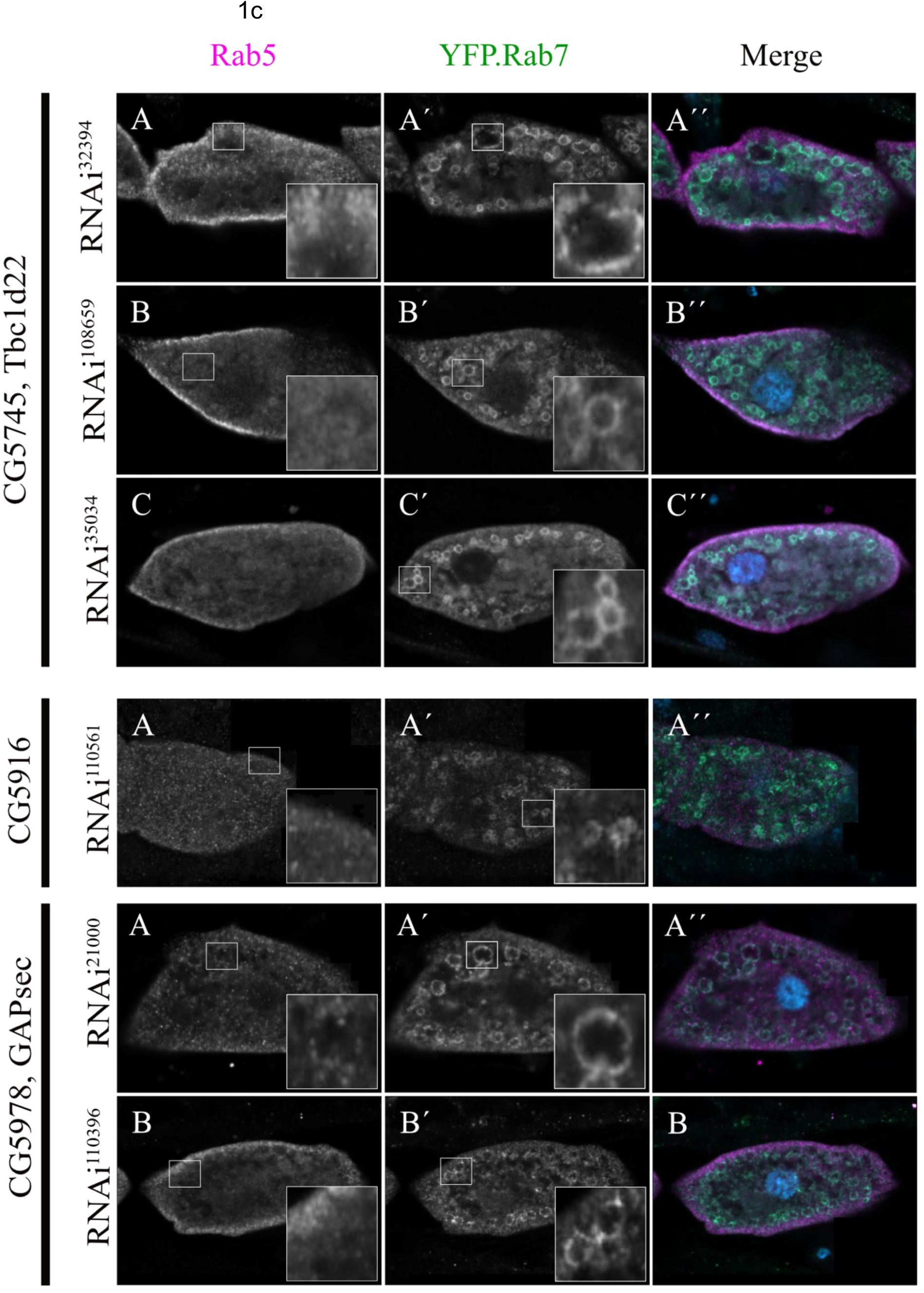
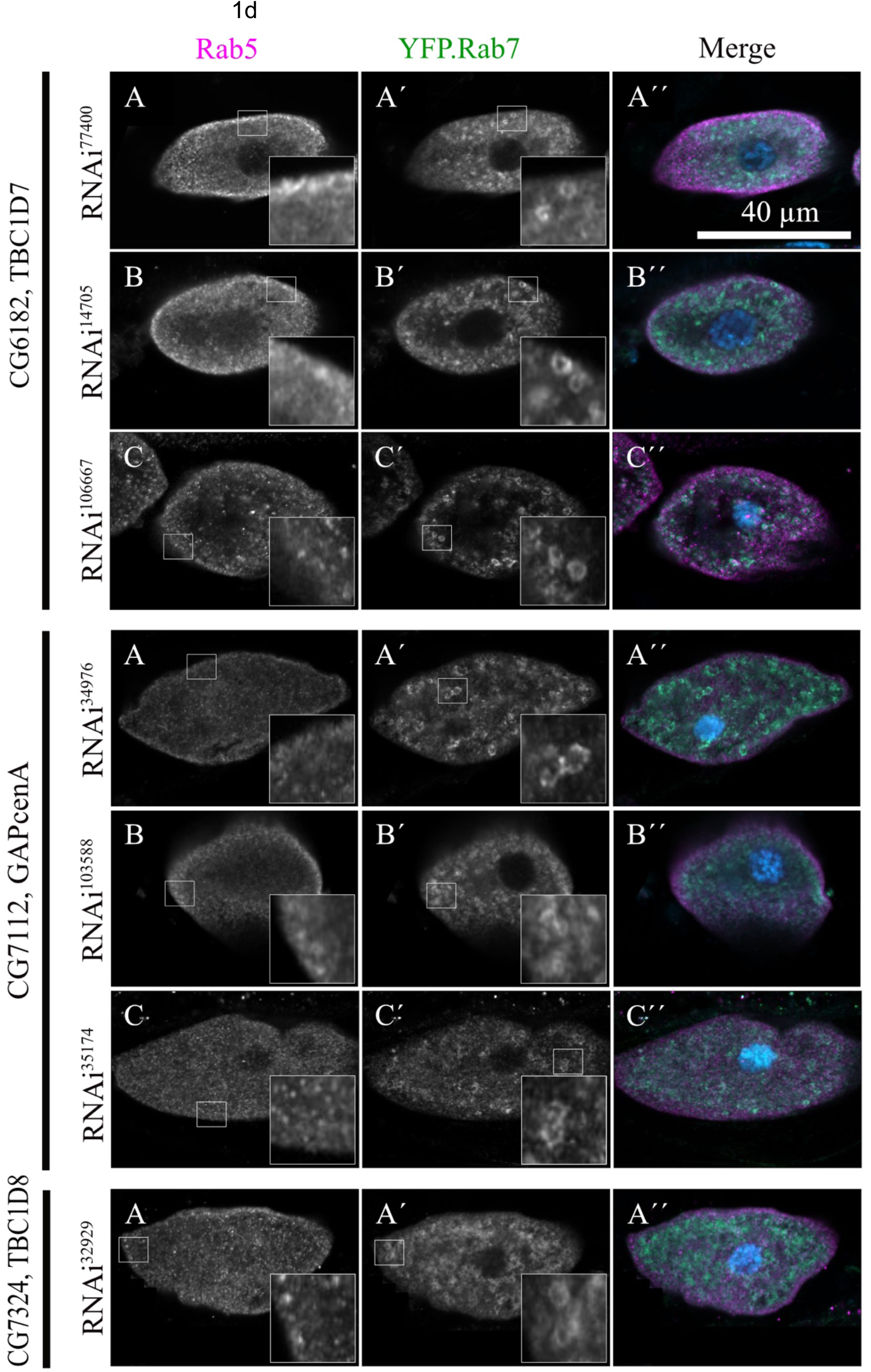
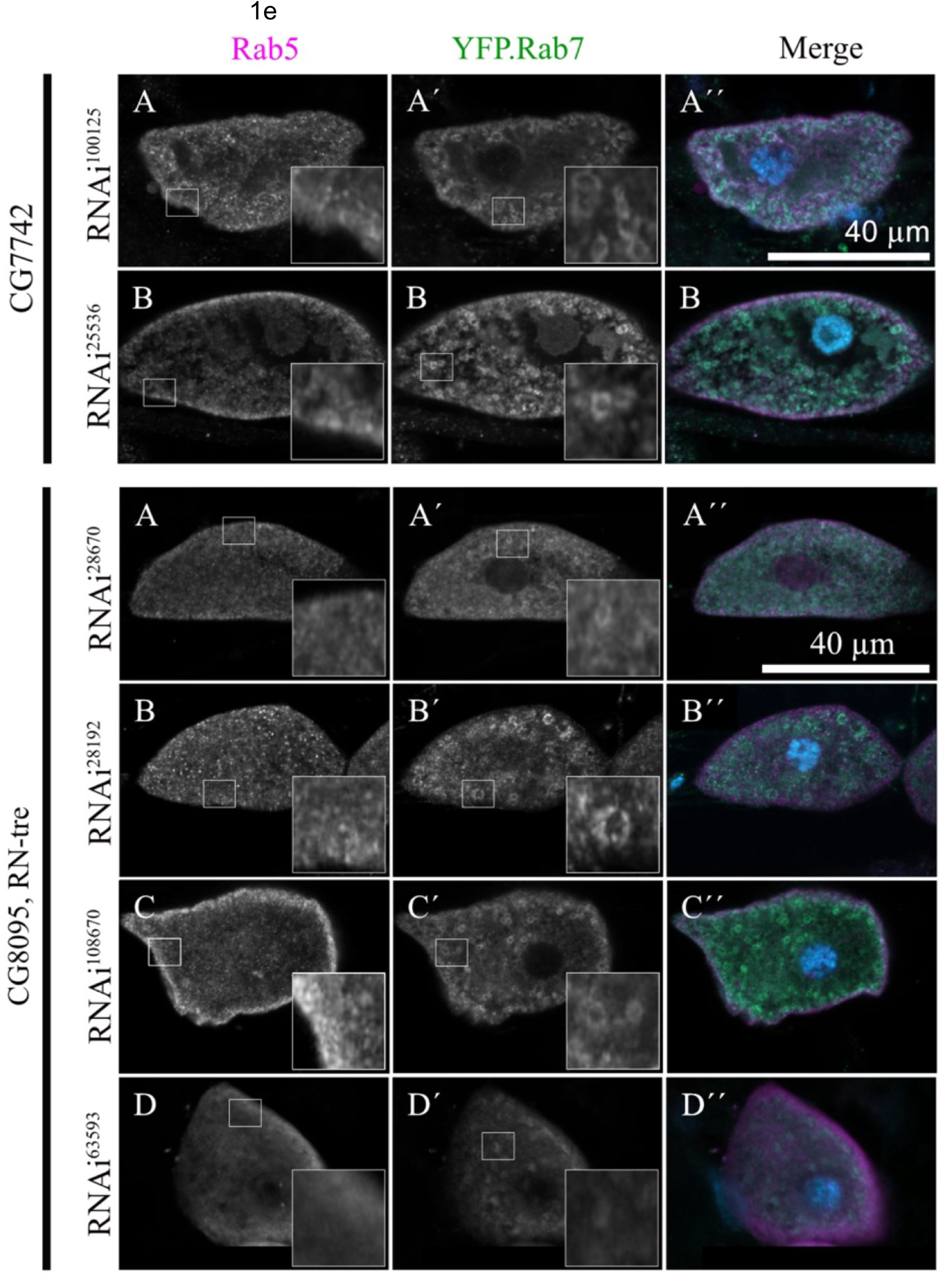
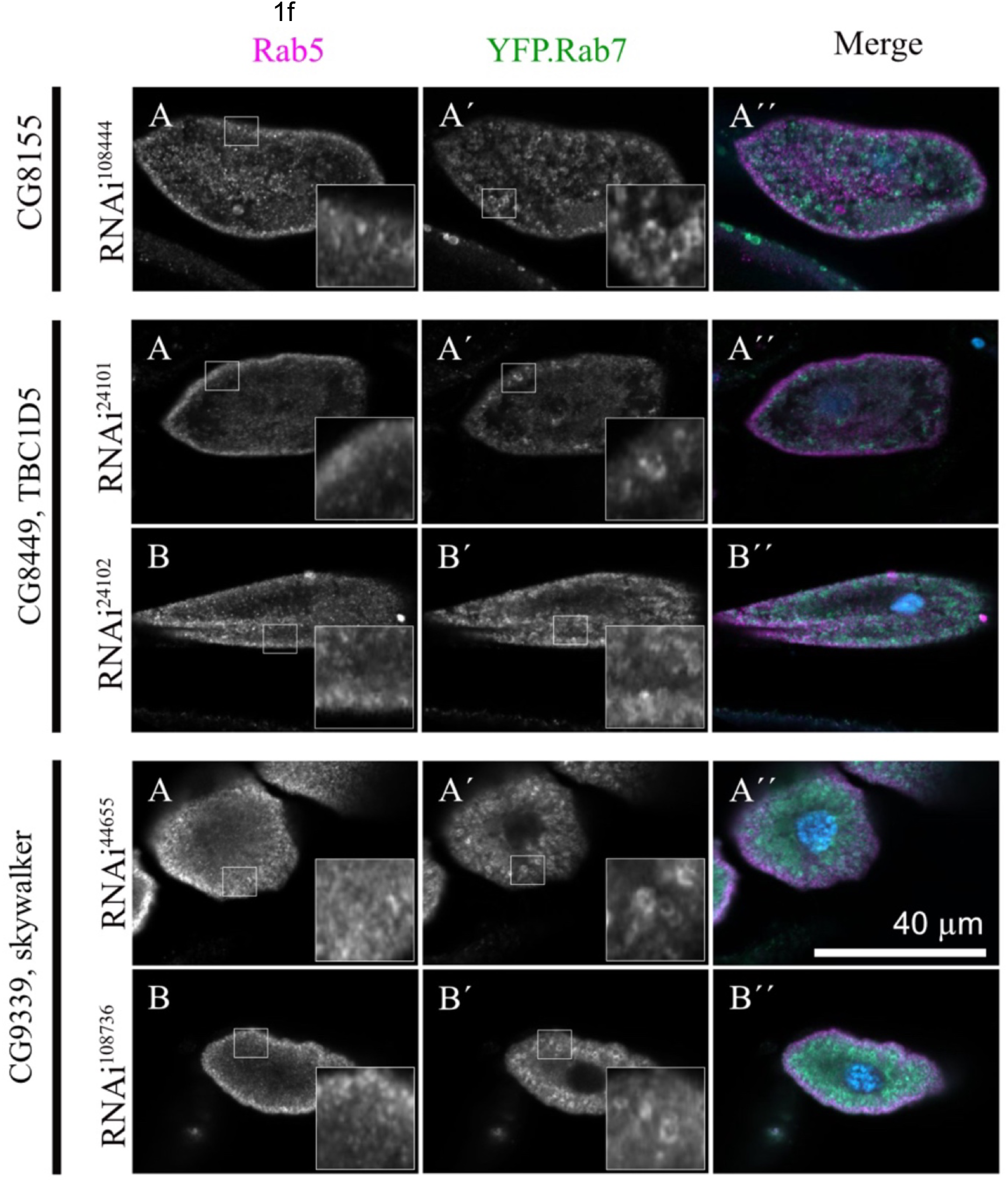
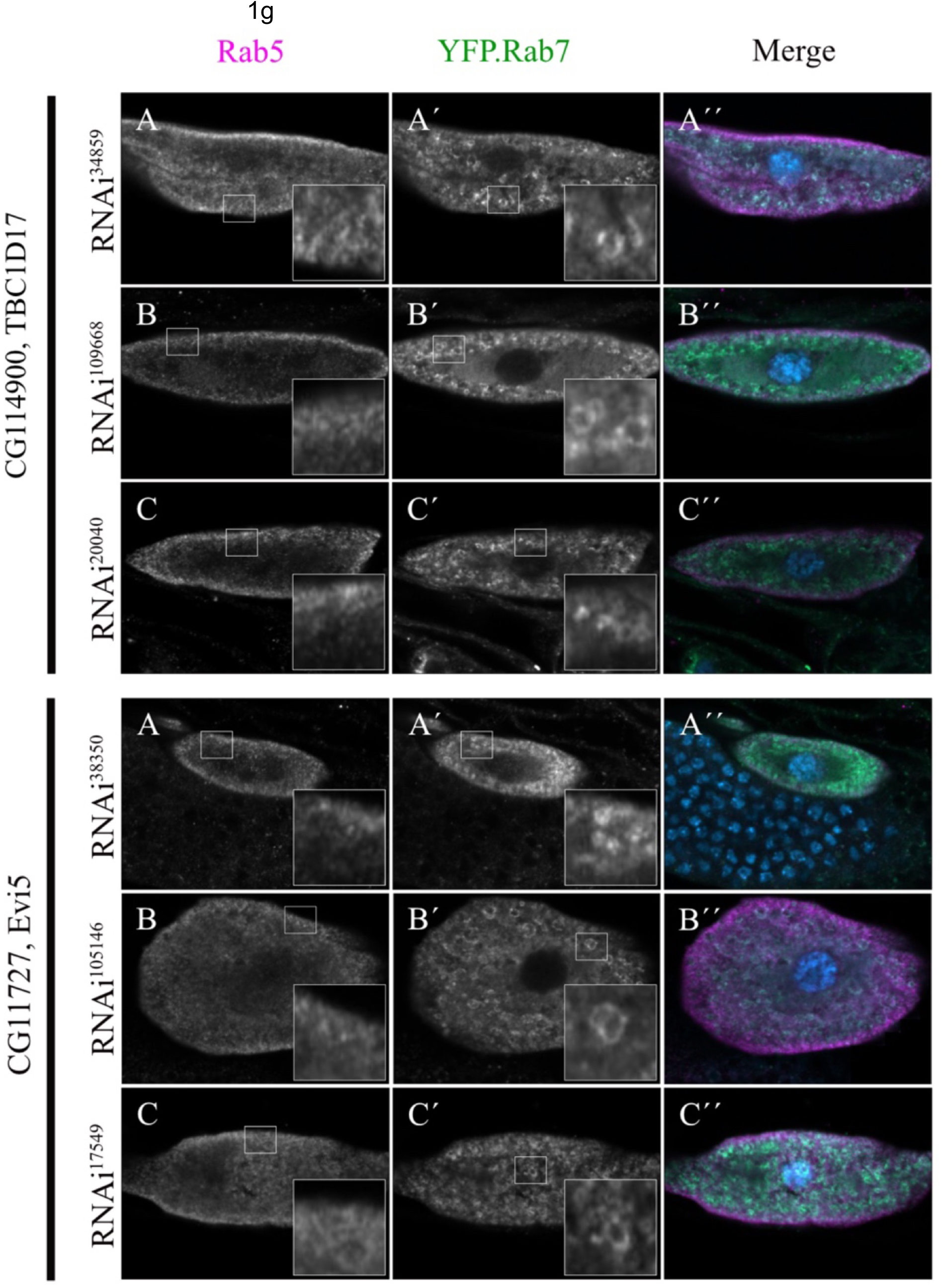
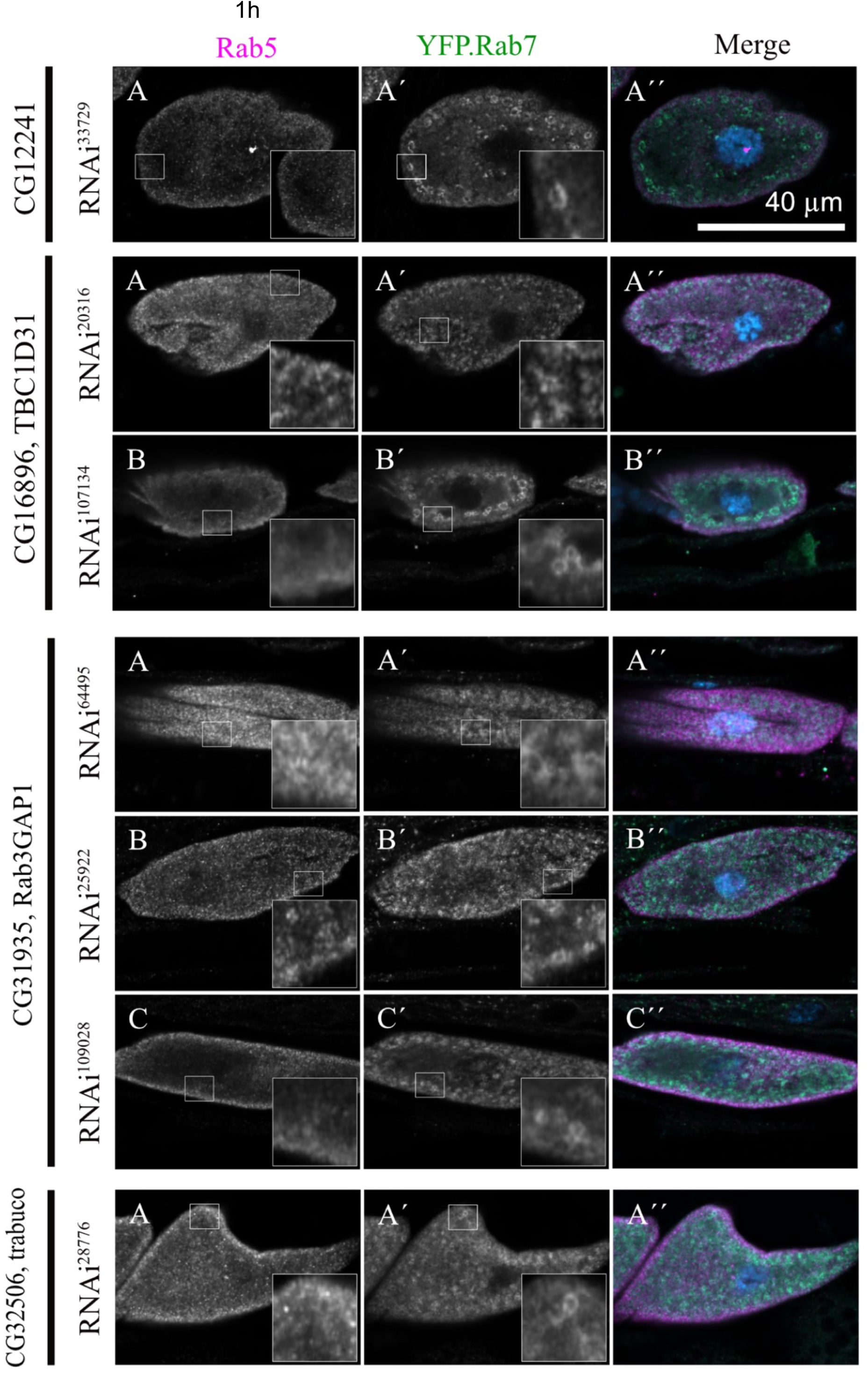
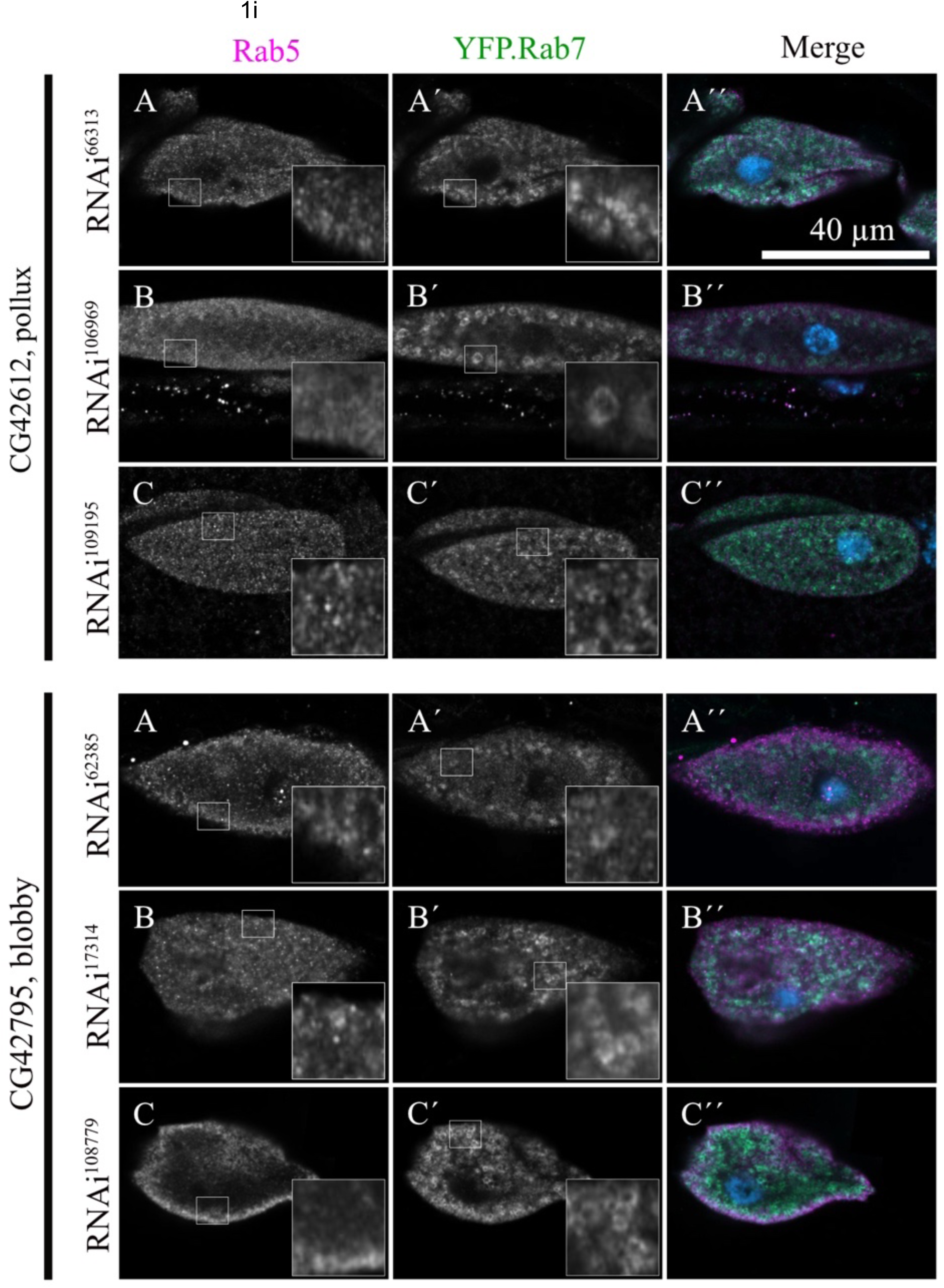
a-i: Distribution of Rab5 and Rab7 in pericardial nephrocytes expressing RNA hairpins for different GAPs. Pericardial nephrocytes from dissected 3^rd^ instar larvae of control flies and of flies, in which one of the potential *Drosophila* GAP genes was down-regulated by available RNAi-constructs, were stained for Rab5 (anti-Rab5) and Rab7 (anti-YFP.Rab7) to identify potential endosomal maturation defects.

**Supplementary Figure 2.**
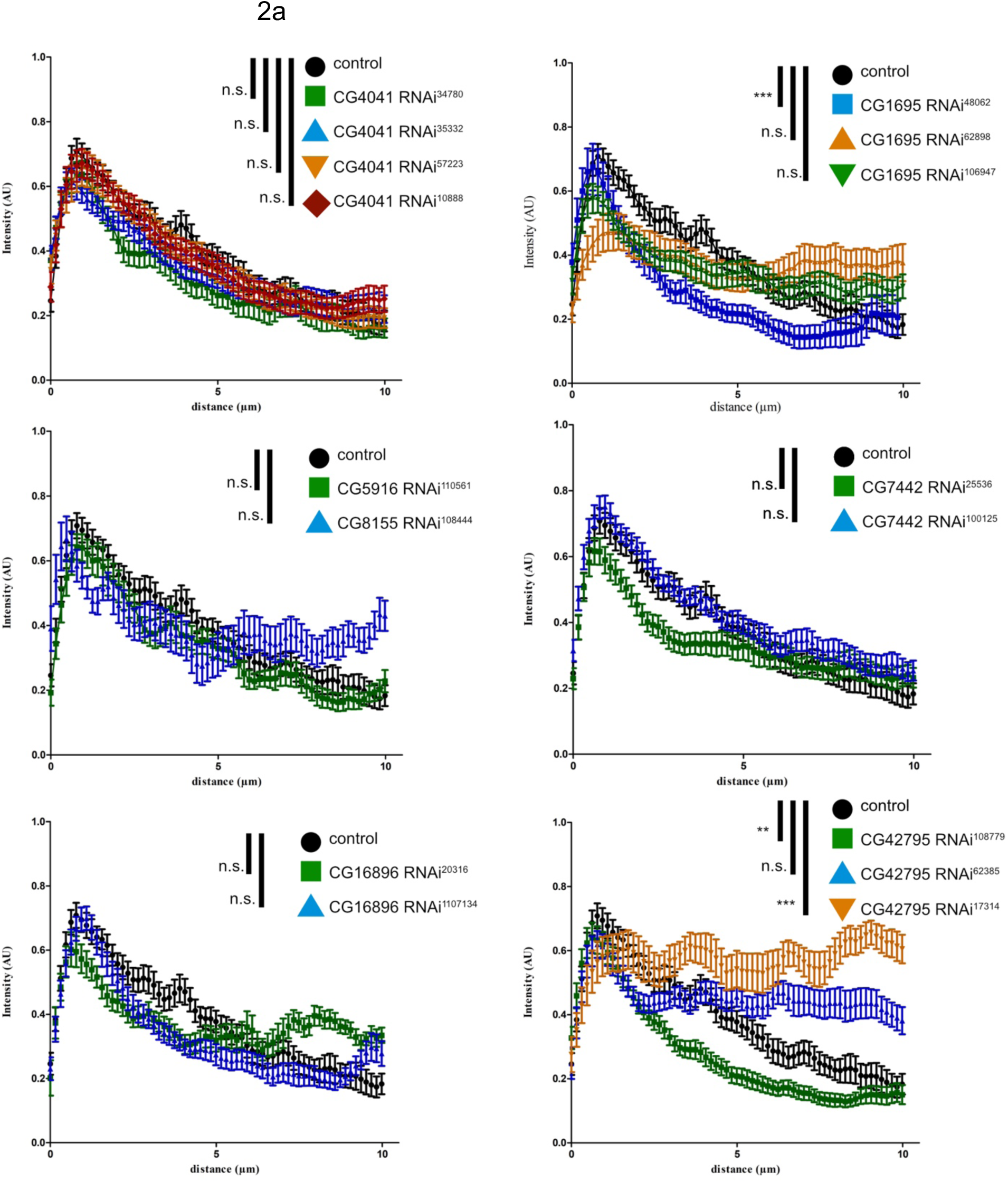
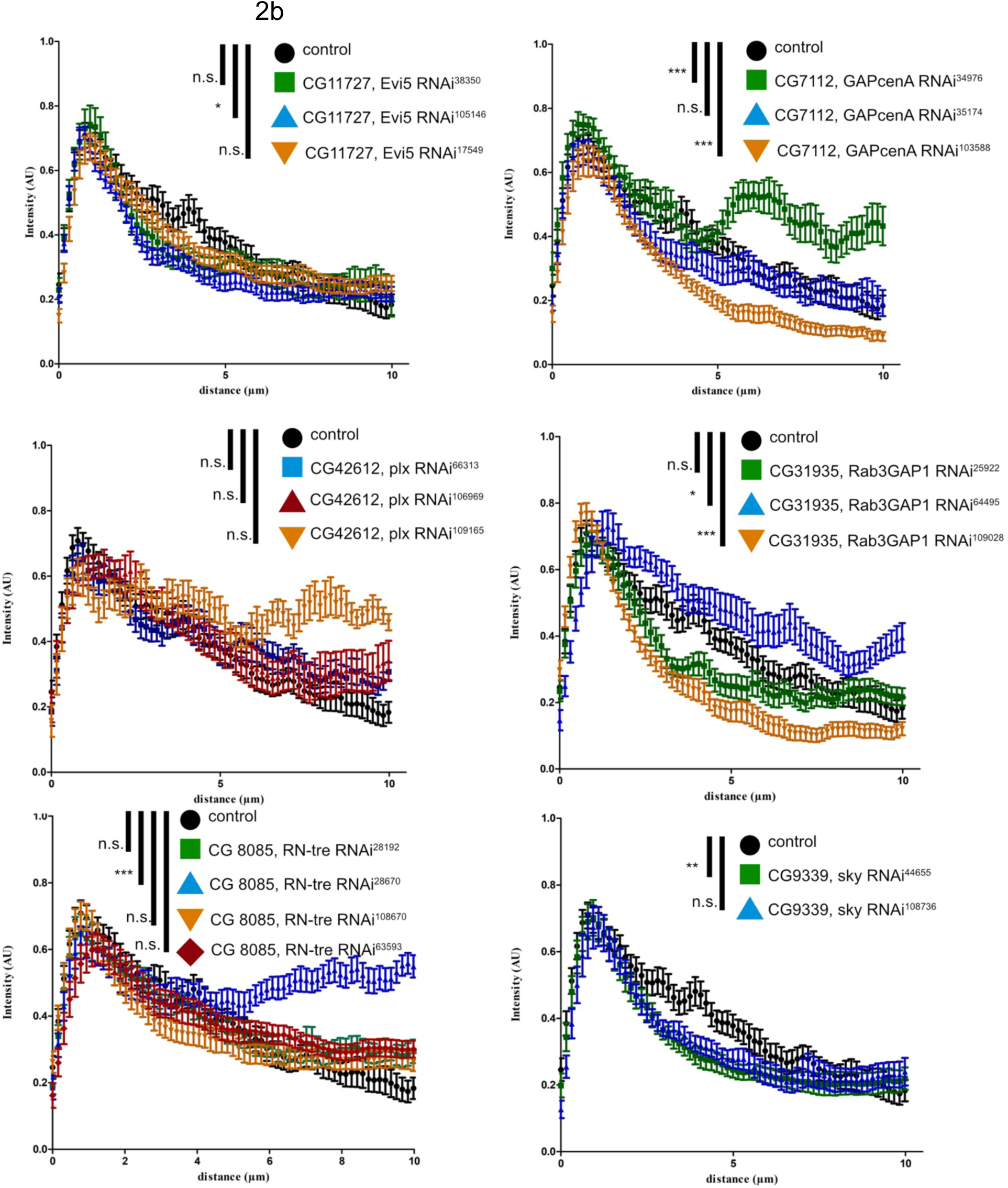
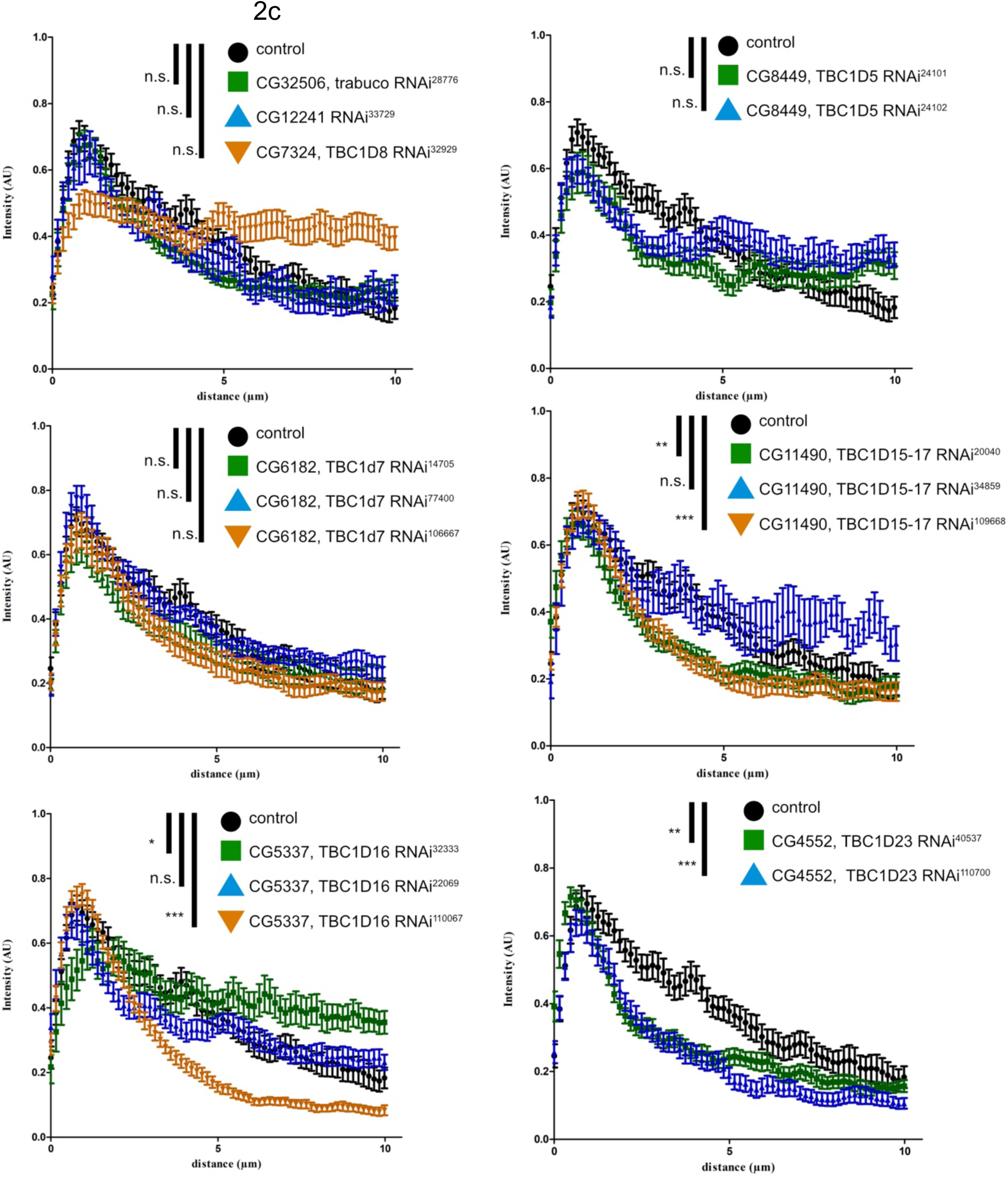
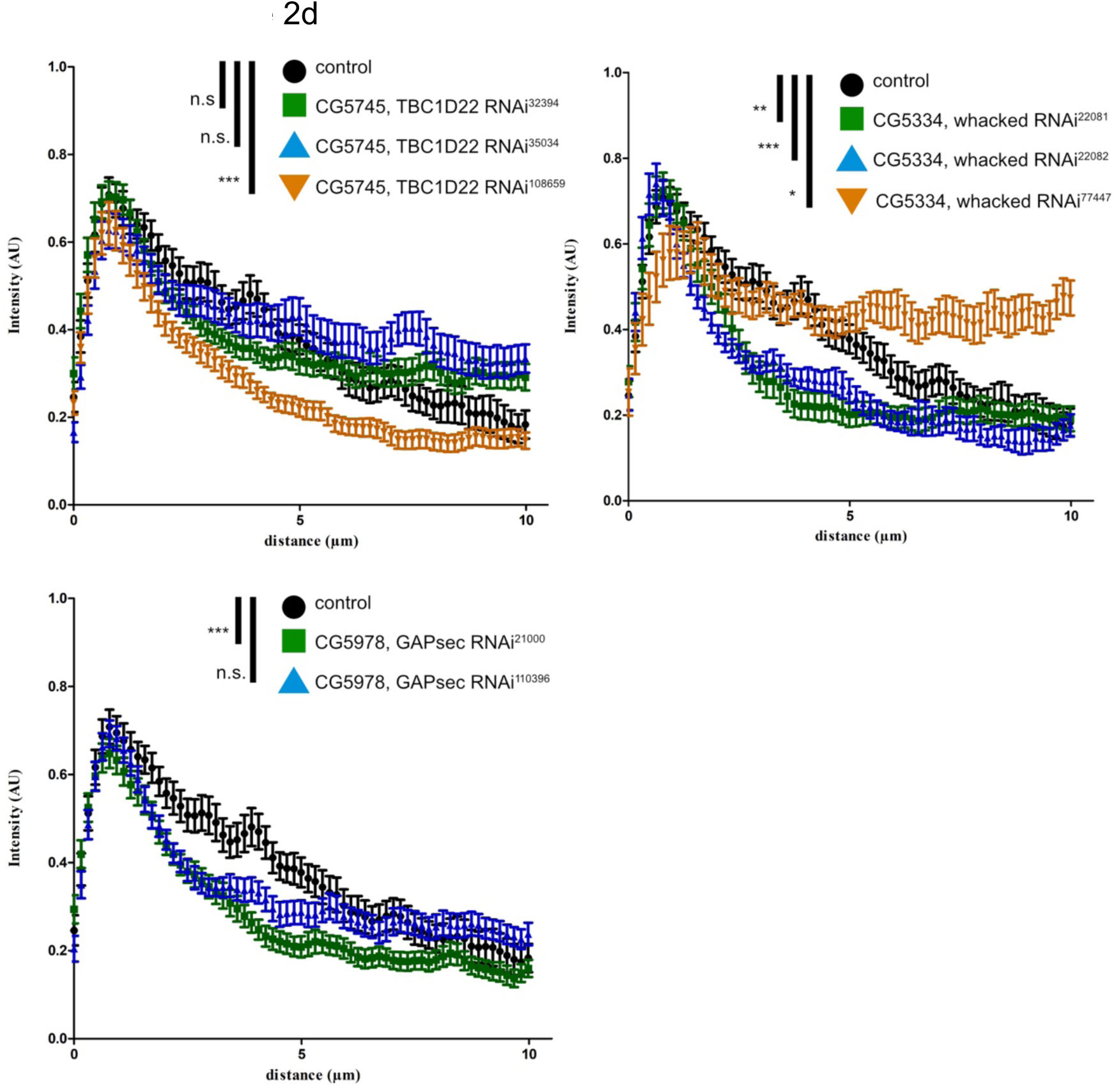
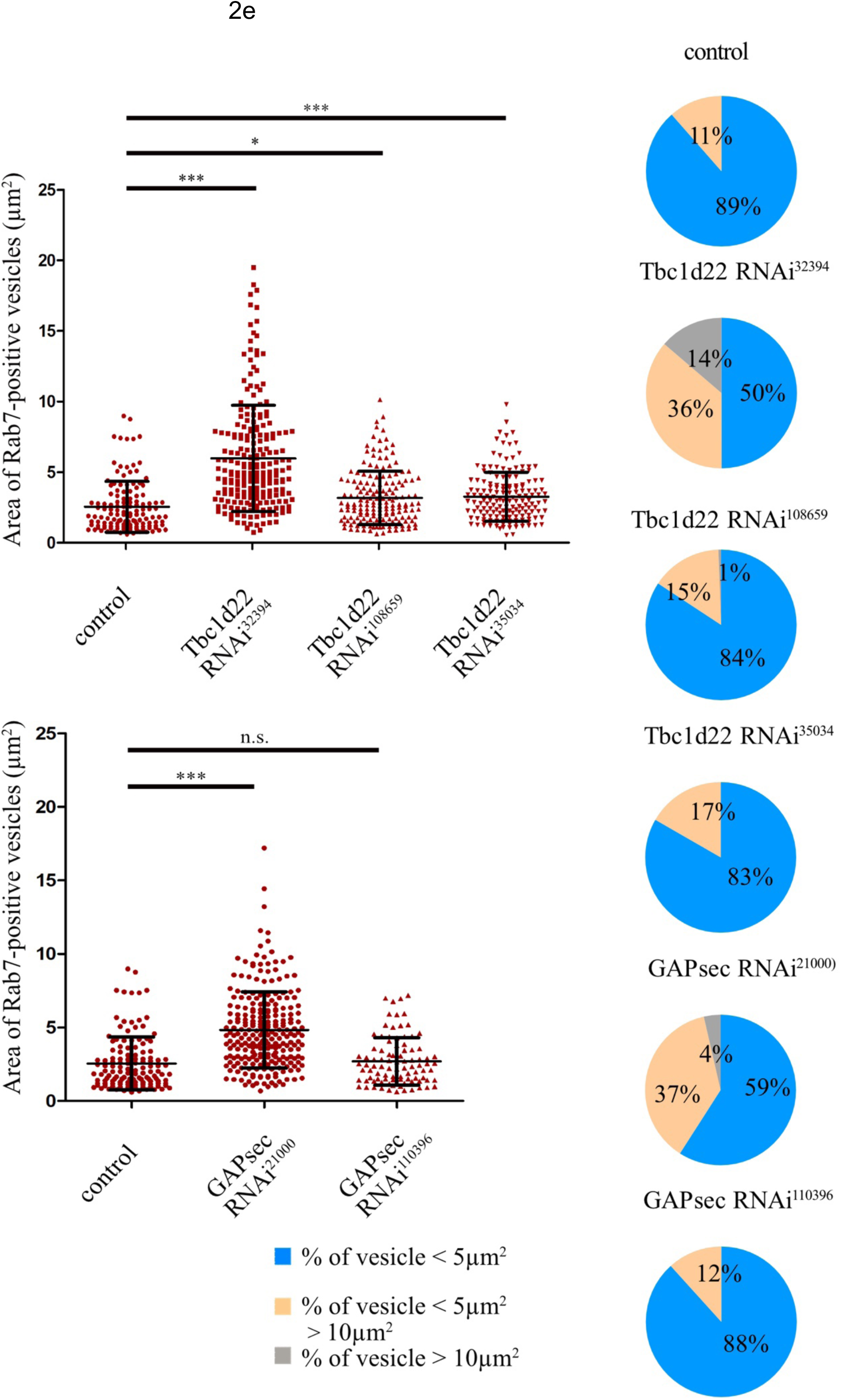
a-e: Quantification of the distribution of Rab5 and Rab7 distribution in pericardial nephrocytes expressing RNA hairpins for different GAPs. Rab5 distribution in nephrocytes along a defined sector of 20 pixels in width and 10 μm in length. The starting point of teaching measurement was the outermost cell periphery. The control was always plotted in black. The “Polygon Selection” tool in ImageJ was used to manually encircle the Rab7-positive vesicles for the area analysis. All measured vesicles were inserted into the ROI Manager, and the area of each vesicle was calculated. Further data processing was performed using Microsoft Excel, and vesicles were subdivided into three groups according to their area: smaller than 5 µm^2^, between 5 µm^2^ and 10 µm^2^ and larger than 10 µm^2^. Prism was used for statistical analysis. The normal distribution was determined using the KS normality test, the D’Agestino & Pearson omnibus normality test, and the Shapiro-Wilk normality test. The total values of the different genotypes were then compared using the Kruskal-Wallis test. Asterisks indicate significance values. * p<0.5, n.s. not significant.

